# Dietary change without caloric restriction maintains a youthful profile in ageing yeast

**DOI:** 10.1101/2022.07.19.500645

**Authors:** Dorottya Horkai, Jonathan Houseley

## Abstract

Caloric restriction increases lifespan and improves ageing health, but it is unknown whether these outcomes can be separated or achieved through less severe interventions. Here we show that an unrestricted galactose diet in early life minimises change during replicative ageing in budding yeast, irrespective of diet later in life. Lifespan and average mother cell division rate are comparable between glucose and galactose diets, but markers of senescence and the progressive dysregulation of gene expression observed on glucose are minimal on galactose, showing these to be associated rather than intrinsic aspects of the replicative ageing process. Respiration on galactose is critical for minimising hallmarks of ageing, and forced respiration during ageing on glucose by over-expression of the mitochondrial biogenesis factor Hap4 also has the same effect though only in a fraction of cells. This fraction maintains Hap4 activity to advanced age with low senescence and a youthful gene expression profile, whereas other cells in the same population lose Hap4 activity, undergo dramatic dysregulation of gene expression and accumulate fragments of chromosome XII (ChrXIIr), which are tightly associated with senescence. Our findings support the existence of two separable ageing trajectories in yeast. We propose that a complete shift to the healthy ageing mode can be achieved in wild-type cells through dietary change in early life without restriction.

## Introduction

Dietary restriction can improve ageing health in eukaryotes ranging from budding yeast to primates and even humans, but remains an unrealistic approach for improving ageing health in human populations as this sacrifice requires substantial commitment (Didier et al., 2016; Fontana and Klein, 2007; Fontana et al., 2010; Green et al., 2022; Hofer et al., 2022). However, the beneficial effects of dietary restriction are not simply due to a reduction in calories as equivalent effects can be achieved through limiting or even tailoring amino acid intake, indicating that improved ageing health results from altered metabolic state rather than minimal energy intake (Jiang et al., 2000; Mair et al., 2005; Trautman et al., 2022). Improved ageing health may therefore be achievable in humans through optimising diet, but this will require a much better understanding of the interplay between diet and the biological mechanisms leading to ageing pathology (Lee et al., 2021).

Replicative ageing of the budding yeast *S. cerevisiae* is routinely employed as a rapid and genetically tractable model system for ageing research. Budding yeast cells divide asymmetrically into a mother cell and a daughter cell, with the mother cell undergoing only a limited number of divisions before permanently exiting the cell cycle (Mortimer and Johnston, 1959). Mother cells maintain a rapid and uniform cell cycle for most of their replicative lifespan but the cell cycle slows dramatically in the last few divisions (Egilmez and Jazwinski, 1989; Fehrmann et al., 2013; Mortimer and Johnston, 1959), a point designated the Senescence Entry Point (SEP) that is coincident with the appearance of apparently pathological markers including mitochondrial protein foci formation and nucleolar expansion (Fehrmann et al., 2013; Morlot et al., 2019). Many other changes occur during replicative ageing, including cell growth (Mortimer and Johnston, 1959; Neurohr et al., 2019), loss of mitochondrial membrane potential (Veatch et al., 2009), mitochondrial fragmentation (Lam et al., 2011), loss of vacuolar acidity (Hughes and Gottschling, 2012), nuclear pore disruption (Meinema et al., 2022), and protein oxidation and aggregation (Aguilaniu et al., 2003; Erjavec et al., 2008), each of which likely contributes to the eventual loss of replicative potential, but how these relate to each other or to the extension of cell cycle timing at the SEP remains largely unknown.

Genetic and epigenetic instability are hallmarks of ageing, and both processes are observed during yeast replicative ageing (Lopez-Otin et al., 2013). Although the yeast genome does not accrue significant mutations with age, recombination events within the ribosomal DNA (rDNA) give rise to extrachromosomal rDNA circles (ERCs), which accumulate massively with age to add 30-40% to total genome size in a single day (Cruz et al., 2018; Kaya et al., 2015; Sinclair and Guarente, 1997). Chromatin structure and the genomic distribution of histone modifications also change dramatically with age, either as cause or consequence of gene expression change (Cruz et al., 2018; Dang et al., 2009; Hu et al., 2014). Progressive dysregulation of gene expression accompanies yeast replicative ageing (Hendrickson et al., 2018; Hu et al., 2014; Janssens et al., 2015; Kamei et al., 2014; Yiu et al., 2008), with consequent remodelling of the proteome and metabolome in a manner consistent with activation of the environmental stress response (Gasch et al., 2000; Janssens et al., 2015; Lesur and Campbell, 2004; Leupold et al., 2019; Yiu et al., 2008). However, it is not known whether these changes are pathogenic or protective, nor is it clear whether gene expression dysregulation is a consequence of genomic change.

Recent studies have revealed heterogeneity in ageing trajectory, with cells following either a slow dividing, short lived trajectory marked by upregulation of iron transporters and accumulation of the Tom70-GFP mitochondrial protein foci linked to SEP onset, or a fast dividing, long lived trajectory, marked by nucleolar expansion and weak rDNA silencing (Jin et al., 2019; Li et al., 2020; Zhang et al., 2020). The association of weak rDNA silencing with a long lived trajectory is surprising as rDNA silencing mutants such as *sir2*Δ undergo higher rates of rDNA recombination and ERC formation due to interference between non-coding RNA (ncRNA) transcription and sister chromatid cohesion (Gottlieb and Esposito, 1989; Kaeberlein et al., 1999; Kobayashi and Ganley, 2005; Saka et al., 2013). Furthermore, other studies find nucleolar expansion to be linked to onset of the SEP, defects in ribosome precursor export and low iron transporter induction (Chen et al., 2020; Morlot et al., 2019). Therefore, the link between mitochondrial dysfunction and senescence is supported across studies, but nucleolar changes are more variable across experimental systems and the importance and timing of rDNA instability remains to be clarified. A major contributor to these differences may be variation in rDNA copy number between strains, which changes during routine yeast transformation and influences both lifespan and rDNA recombination rate (Hotz et al., 2022; Kwan et al., 2016).

Dietary restriction is most commonly achieved in yeast through caloric restriction - a reduction in media glucose concentration from 2% to 0.5% that reproducibly increases lifespan (Lin et al., 2000). This has been ascribed both to a reduction in growth signalling through PKA and TOR pathways (Kaeberlein et al., 2005; Lin et al., 2000), and an induction of respiration acting through Sir2 to improve rDNA stability and decrease ERC formation (Anderson et al., 2003; Lin et al., 2004; Lin et al., 2002; Schleit et al., 2013). Unfortunately, it is technically very difficult to purify yeast aged under caloric restriction so the long-term physiological effects of caloric restriction remain largely unexplored. Nonetheless, cells aged under caloric restriction do show increased cell cycle time at the end of replicative lifespan, suggesting that the SEP is not repressed (Jo et al., 2015). Unexpectedly, we have noted that changing the primary carbon source during yeast ageing improves the homogeneity of cycling times in aged cells, suggesting that ageing health is improved (Frenk et al., 2017). Here, we demonstrate that growth on galactose as an alternate carbon source in early life dramatically improves ageing health of yeast through enhanced respiration without increasing lifespan, and suppresses both gene expression dysregulation and genome instability.

## Results

### Suppression of age-linked phenotypic change on an unrestricted galactose diet

*S. cerevisiae* preferentially metabolise glucose, but the fermentable sugars galactose and fructose also support robust growth in culture. To determine the effect of non-preferred carbon sources on ageing, we used an optimised Mother Enrichment Program (MEP) protocol in which cells are biotin labelled then aged in rich media, fixed and mother cells purified purified (Figure 1A) (Cruz et al., 2018; Lindstrom and Gottschling, 2009). The MEP system renders daughter cells inviable, such that mother cells can reach very advanced age without media components being exhausted by proliferation of offspring.

**Figure 1:**
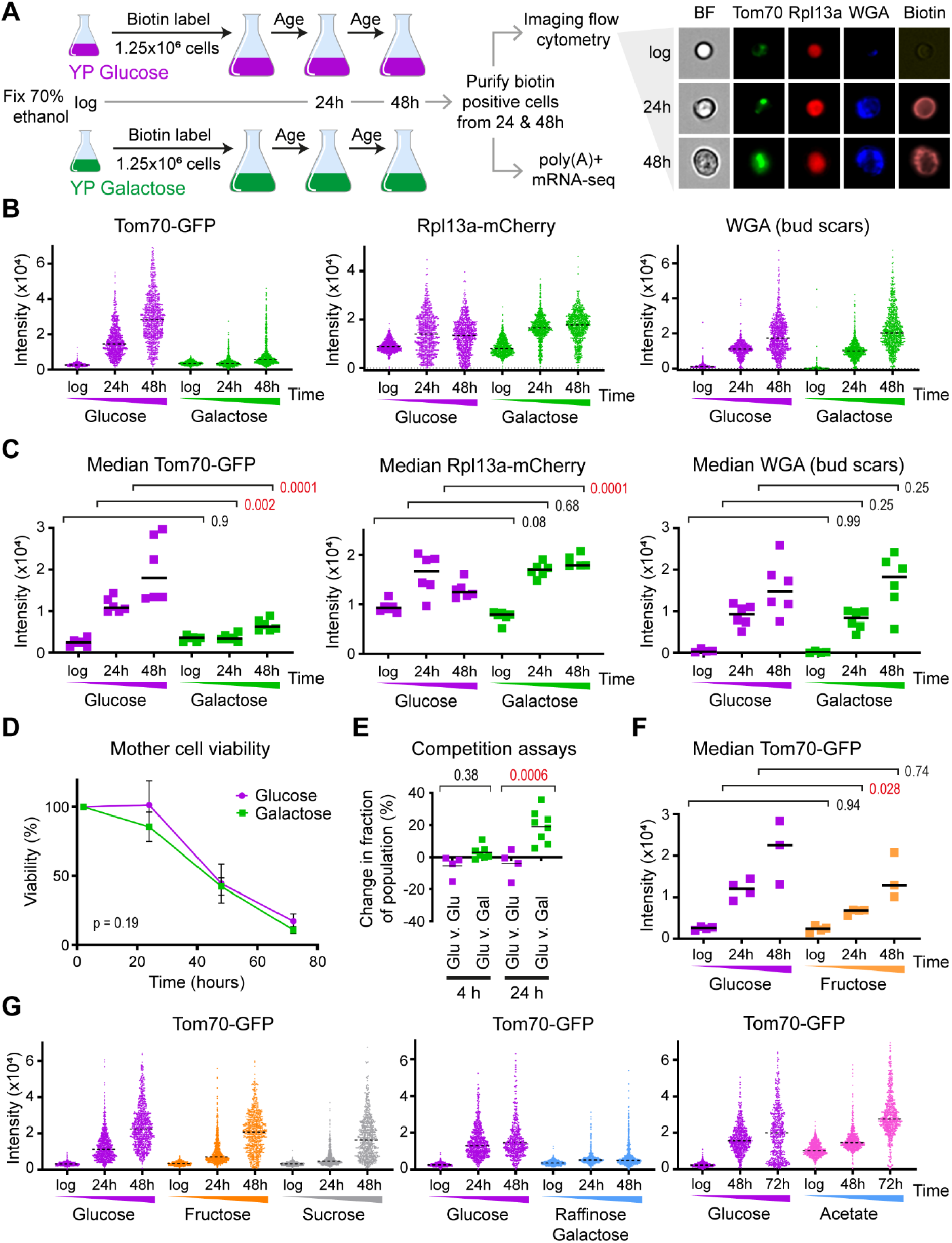
Progression of age-linked phenotypes on different diets. A: Schematic of ageing cell culture and analysis, with examples of imaging flow cytometry data (Tom70-GFP, Rpl13a-mCherry, WGA-Alexa405 and brightfield) for wild type MEP cells ageing on YPD (rich glucose media). B: Distribution of signal intensities from flow cytometry images of Tom70-GFP, Rpl13a-mCherry and WGA-Alexa405 in populations of log phase cells or mother cells aged for 24 and 48 hours in YP with 2% glucose or galactose. 1000 cells are imaged per population after gating for circularity (all) and biotin (aged cells only, based on streptavidin-Alexa647). Images are post-filtered for focus of Tom70-GFP, leaving ∼600 cells per population. C: Median signal intensities from multiple biological replicates of populations described in B. n=6, p values calculated by 2 way ANOVA with post-hoc Tukey test for age and diet. D: Lifespan measured in YPD and YPGal media based on %age of viable cells remaining at each time point. p value from t test comparing area under curves, n=5 for glucose and for galactose. E: Competition assays measuring fitness in glucose between cells aged for indicated times on glucose or galactose. Wild-type MEP cells carrying G418 or Nourseothricin resistance genes were aged for 4 and for 24 hours post estradiol addition in YP media with 2% glucose or galactose, then mixed in pairs carrying reciprocal markers for competition and plated on G418 and on nourseothricin plates at t=0 to determine initial mixture competition, then inoculated at 1:1,000 in YPD and allowed to grow to saturation before plating again on G418 and on nourseothricin plates. Change in composition (%) from t=0 to end of competition is plotted on y axis. n=4 glucose v glucose, n=8 glucose v galactose, p values calculated by 1 way ANOVA with post-hoc Tukey test. F: Median Tom70-GFP intensity for populations aged on glucose or fructose, n=4 (log, 24h) n=3 (48h), analysed as in C. Matching Rpl13a-mCherry and WGA data shown in Figure S1C. G: Population distributions of Tom70-GFP for cells aged on indicated carbon sources, matching Rpl13a-mCherry and WGA data shown in Figures S1D and S1E. Note extended time-course glucose control for acetate experiment, however 72 hour acetate is age-matched to 48 hour glucose as cell division is slower on acetate.

The Senescence Entry Point (SEP) represents an abrupt transition in ageing at which yeast cells cease to divide rapidly and the cell cycle becomes slow and heterogeneous (Fehrmann et al., 2013). Post-SEP cells are marked by highly fluorescent foci of the GFP-tagged mitochondrial protein Tom70 in G1 (Fehrmann et al., 2013; Li et al., 2020). To quantify Tom70-GFP fluorescence as a SEP marker, we optimised an imaging flow cytometry assay for aged MEP cells expressing Tom70-GFP as well as an Rpl13a-mCherry marker that accumulates only slightly with age (Janssens and Veenhoff, 2016). Per sample, 1000 purified mother cells gated for streptavidin and roundness are imaged for brightfield, Tom70-GFP, Rpl13a-mCherry and WGA-AF405 to label bud scars (Figure 1A right).

Tom70-GFP increased substantially with age across the whole population in glucose (Figure 1B left, glucose), which was expected as the roundness gate selects towards G1 cells in which Tom70-GFP foci manifest (Fehrmann et al., 2013). In contrast, Rpl13a-mCherry increased slightly but became much more heterogeneous at 24 hours and often declined from 24 to 48 hours, perhaps reflecting the onset of ribophagy once cells cease to divide (Figure 1B middle, glucose). WGA fluorescence increased progressively and correlated to the accumulation of bud scars on both glucose and galactose (Figure S1A), forming a useful metric for replicative age as previously noted (Chen and Contreras, 2007; Janssens et al., 2015; Patterson and Maxwell, 2014). Imaging flow cytometry therefore allows rapid quantification of age and age-associated phenotypes, with excellent reproducibility across biological replicates (Figure 1C).

Remarkably, when glucose was replaced with galactose, the Tom70-GFP and Rpl13a-mCherry markers behaved very differently: Tom70-GFP increased only marginally with age (Figures 1B & 1C left), whilst the Rpl13a-mCherry signal increased at 24 hours and remained high from 24 to 48 hours without becoming heterogeneous (Figures 1B & 1C middle). These differences suggest that ageing of MEP diploid cells in galactose is not associated with a SEP, despite cells reaching the same replicative age after 24 and 48 hours as measured by both WGA fluorescence intensity and manual bud scar counting (Figure 1C right and Figure S1A). Mitochondrial dysfunction in aged cells is associated with defects in the expansion of the vacuole (Aufschnaiter and Buttner, 2019; Hughes and Gottschling, 2012), so we examined the vacuolar marker Vph1-mCherry to reinforce the mitochondrial Tom70-GFP phenotype. The Vph1-mCherry signal also increased substantially with age in glucose, showing that the Tom70-GFP phenotype is not an artefact of a single protein or fluorescent tag, but was almost unchanged with age in galactose (Figure S1B). Therefore, ageing phenotypes revealed by multiple markers are suppressed when cells are aged on galactose instead of glucose.

The glucose-galactose differences could reflect a dramatic extension of lifespan such that the SEP is simply delayed, but the viability of cells on glucose and galactose decayed at a similar rate over time in culture (Figure 1D). Given that bud scar counts and WGA intensity at 24 and 48 hours are equivalent lifespan cannot be substantially different between glucose and galactose (Figure 1C and S1A), in accord with a previous report that lifespan is slightly shorter on galactose (Liu et al., 2015). Alternatively, these markers may not accurately report cellular fitness, however when directly competed in glucose media, cells aged in galactose out-perform cells equivalently aged in glucose (Figure 1E). This means that despite equivalent lifespan, diploid MEP cells aged on galactose do not show a SEP and remain fitter than those aged on glucose.

In contrast, fructose did not strongly suppress ageing phenotypes: Tom70-GFP was somewhat reduced at 24 hours and 48 hours relative to glucose ageing but was still higher than at log phase and the difference was only significant at 24 hours (Figure 1F). Furthermore, the behaviour of the Rpl13a-mCherry marker was indistinguishable between glucose and fructose (Figure S1C). This indicated that different diets have different effects on ageing in yeast so we surveyed other carbon sources (Figure 1G, Figure S1D & S1E): sucrose acted very similarly to fructose, a 2% raffinose 0.5% galactose mixture was similar to galactose alone, whilst acetate was exceptional as Tom70-GFP accumulated similarly to cells in glucose but Rpl13a-mCherry did not decrease at advanced ages. Acetate is likely to be exceptional anyway as cell division is very slow (hence the reduced WGA signal at each time compared to glucose) and mitochondrial biogenesis is high, which will contribute to the Tom70-GFP signal independent of the SEP.

Taken together, these experiments reveal that cells aged in galactose undergo minimal age-linked changes based on multiple fluorescence markers of the SEP and direct measurement of fitness. This occurs without an increase in lifespan, proving that at least in yeast lifespan is distinct from senescence and loss of fitness, and must represent a separable coincident process.

### The transcriptome undergoes minimal dysregulation during ageing in galactose

Many studies have reported characteristic changes in gene expression pattern during yeast replicative ageing on glucose (Cruz et al., 2018; Hendrickson et al., 2018; Hu et al., 2014; Janssens et al., 2015; Kamei et al., 2014; Yiu et al., 2008). This may represent a response to metabolic stresses arising during ageing (for example activation of the Environmental Stress Response), dysregulation of transcription, or both. The dramatic differences in ageing phenotype between glucose and other carbon sources led us to examine the behaviour of the transcriptome, both as an orthogonal marker of the ageing process and to provide mechanistic insights.

Distortion of global gene expression is easily visualised on an MA plot, which for each mRNA plots the change in abundance between two conditions against average abundance. Comparing young and old cells in glucose shows that genes with low average expression undergo large increases in abundance with age, whereas highly expressed genes (mostly ribosome and ribosome biogenesis proteins) are slightly reduced relative to average, resulting in the distribution becoming skewed away from the x-axis in accord with previous studies (Figure 2A left) (Cruz et al., 2018; Hendrickson et al., 2018; Hu et al., 2014; Janssens et al., 2015; Kamei et al., 2014). However, comparison of young and old cells in galactose reveals gene expression changes but only a weak bias towards induction of normally inactive genes between log and 48 hours (Figure 2A right), showing that age-linked dysregulation of mRNA abundance is not an intrinsic aspect of the ageing process.

**Figure 2:**
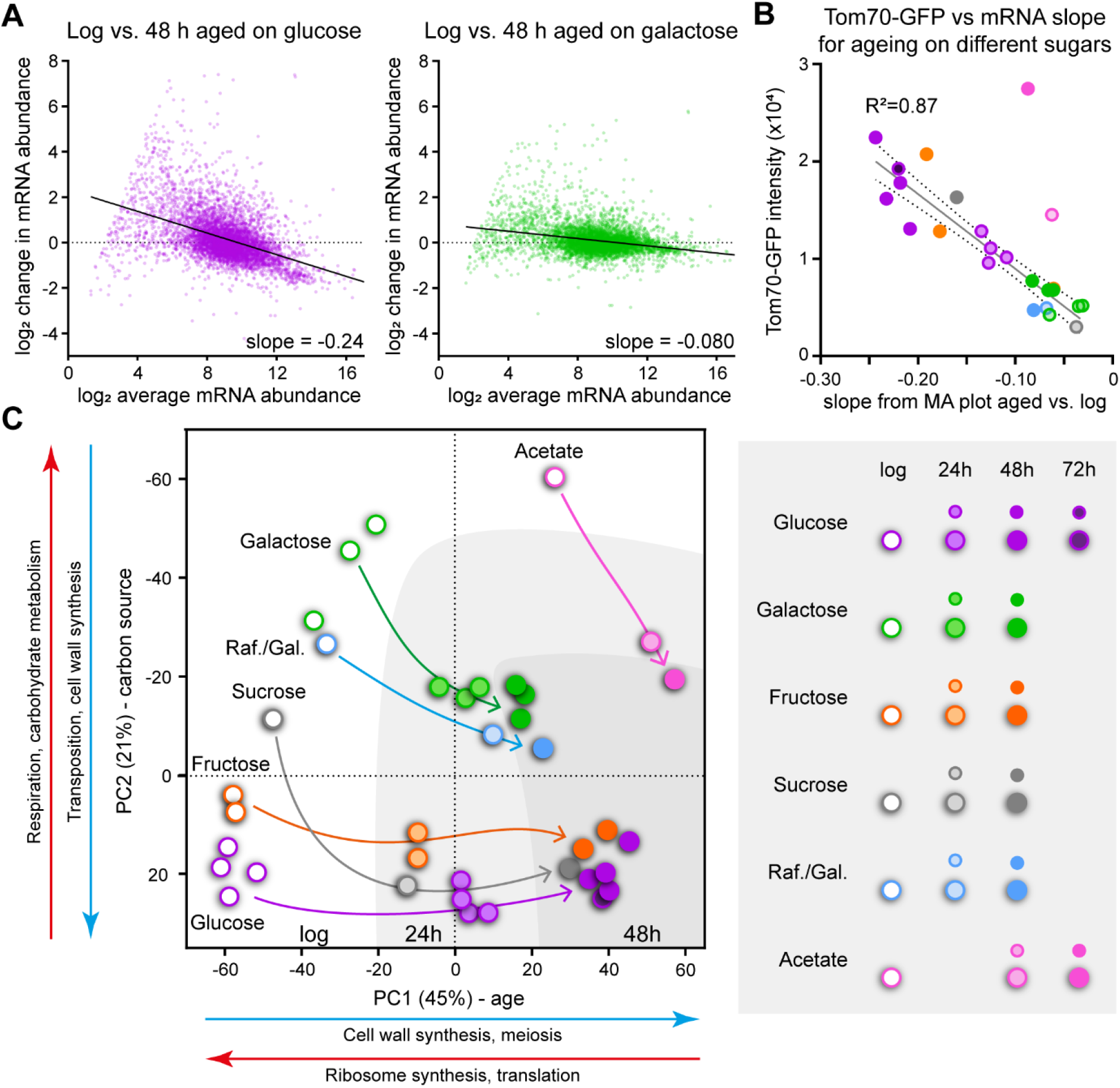
Effects of diet on gene expression change during ageing. A: MA plots showing change in abundance between log and 48 hours relative to average abundance for each mRNA, for cells growing on glucose (left) or galactose (right). Positive values on y-axis indicate increased abundance with age relative to average, negative values indicate decreased abundance, zero indicates no change. Abundances are size-factor normalised log_2_ transformed read counts from poly(A) selected sequencing libraries, genes with <4 counts at log phase were filtered as changes cannot be accurately quantified for these. Data is average of 2 biological replicates, slopes calculated by linear regression. B: Plot of Tom70-GFP intensity against slope values of mRNA distribution change for multiple biological replicate samples of wild type cells aged on different carbon sources. For each individual aged sample, half the cells were used for RNAseq and slope calculated by linear regression from an MA plot of gene expression versus matched log phase sample. Tom70-GFP intensity was obtained from imaging flow cytometry performed on the other half of the sample. Line of best fit calculated by linear regression showing 95% confidence window, acetate samples were excluded as they clearly deviate from the trend. Key to carbon source and age given below graph. C: PCA plot for individual biological replicates at log, 24 and 48 hours of age on different carbon sources. Calculated from size-factor normalised log_2_ transformed read counts for all genes excluding mitochondrial genome-encoded mRNAs. Major GO categories for each PC are indicated based on the 300 highest rotation genes underlying the PC in either direction, full GO analysis in Supplementary File 1.

Transcriptomes of cells aged on fructose, sucrose, raffinose/galactose and acetate show varying degrees of dysregulation (Figure S2) that we captured in a single value - the slope of a line of best fit drawn from an MA plot comparing young and old. These lines (on Figure 2A and Figure S2) have a slope approaching zero in galactose where there is little directional bias in the gene expression changes during ageing, but become increasingly negative with age-linked gene expression dysregulation. This slope value correlates very well to Tom70-GFP intensity across different ages and media conditions (Figure 2B), indicating that transcriptional dysregulation is associated with the onset of the SEP. The only exception is acetate, which has a low slope but a high Tom70-GFP signal that likely reflects very high mitochondrial biogenesis on this strict oxidative phosphorylation substrate rather than SEP-associated Tom70 accumulation.

Principal component analysis (PCA) of this transcriptome data from cells aged on different carbon sources revealed that samples separated by replicative age on the first principal component (PC1, explaining 45% of the dataset variance), and largely by carbon source on PC2 (explaining 21% of variance) (Figure 2C). Glucose and fructose, which are metabolised similarly, cluster together at log and 48 hour timepoints, with fructose 24 hour lagging slightly behind glucose 24 hour on the same trajectory (Figure 2C purple and orange). The transcriptional profile associated with sucrose fermentation differs from glucose and fructose and as such the log phase sucrose profile is separated on PC2 (Marques et al., 2016), however, sucrose is processed into glucose and fructose prior to being metabolised, and aged sucrose transcriptomes are very similar to fructose (Figure 2C grey). Galactose-grown samples migrate less distance on PC1 with age, are separated from glucose and fructose on PC2 at all time points, and change little between 24 hours and 48 hours showing that gene expression is relatively stable (Figure 2C green), with the raffinose/galactose mixture following the same trajectory (Figure 2C blue). Acetate is again the exception as it also moves little on PC1 with age but even at log phase is already coincident on PC1 with highly aged samples from other carbon sources, probably reflecting very slow growth rate (Figure 2C pink). PC1 loading is dominated by increasing cell wall synthesis and decreasing ribosome synthesis, whilst PC2 loading is driven by increasing respiration and decreasing retrotransposition.

These experiments show that the onset of the SEP, as marked by Tom70-GFP foci formation, is tightly linked to gene expression dysregulation on fermentable carbon sources. Given that lifespan is not different between glucose and galactose, this means that gene expression dysregulation is an aspect of ageing that is not intrinsically associated with increasing replicative age or lifespan.

### Respiration protects against the SEP

This RNA-seq dataset should contain mechanistic information about healthy ageing, but the aged datasets are hard to compare between media as global differences in mRNA distribution violate a basic assumption of packages such as DESeq2. We therefore performed a quantile normalisation to remove the global differences before applying DESeq2 to discover recurrent gene expression differences between samples. Pairwise normalised glucose versus galactose comparisons at each age yielded variable gene sets, so we asked the more conservative question of which genes are consistently differentially expressed across all age comparisons (Figure 3A and Supplementary File 1). This much smaller set is enriched for respiration, which is at first sight surprising as galactose and glucose are both fermentable carbon sources. However, whereas respiration is repressed in glucose, it does occur on galactose and is in fact required for normal growth of S288-derived MEP strains (Fukuhara, 2003; Mortimer and Johnston, 1986; Perez-Samper et al., 2018).

**Figure 3:**
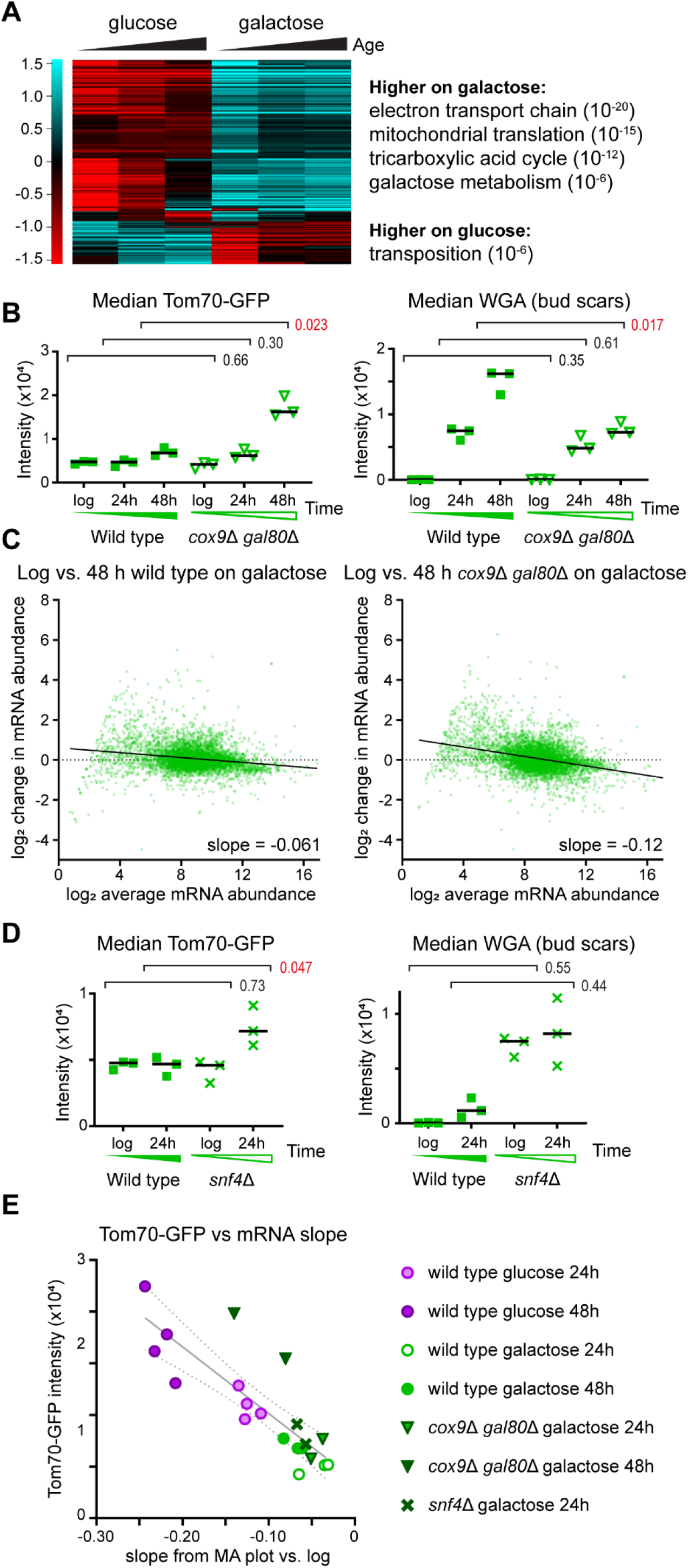
Respiration is required to avoid the SEP and gene expression dysregulation on galactose. A: Hierarchical cluster analysis of genes significantly (p<0.01 by DESeq2 on quantile normalised data) differentially expressed between glucose and galactose at all time points. Average of 2 biological replicates. Principle GO categories with p values are given for each cluster, full GO analysis in Supplementary File 1. B: Imaging flow cytometry for Tom70-GFP and WGA of wild-type and *cox9*Δ *gal80*Δ populations aged on galactose. Performed and analysed as in Figure 1C, n=3. Rpl13a-mCherry data is given in Figure S3A. C: MA plots comparing log and 48 hour-aged populations of wild type and *cox9*Δ *gal80*Δ on galactose, performed as in Figure 2A, n=1 for wild type (matched biological replicate to mutant), n=2 *cox9*Δ *gal80*Δ. D: Imaging flow cytometry for Tom70-GFP and WGA of wild-type and *snf4*Δ populations aged on galactose. Performed and analysed as in Figure 1C, n=3. Rpl13a-mCherry data is given in Figure S3A. E: Plot of Tom70-GFP intensity against slope values of mRNA distribution change, as Figure 2B.

Blocking the electron transport chain to prevent respiration, for example by deleting *COX9*, impairs growth in galactose but this defect can be offset by removing the transcriptional repressor Gal80 (Quarterman et al., 2016). Unlike wild-type, *cox9*Δ *gal80*Δ cells ageing in galactose accumulated substantial Tom70-GFP at 48 hours, despite reaching a much lower replicative age (Figure 3B, Figure S3A), and also underwent greater gene expression dysregulation (Figure 3C). The gene expression phenotype was weaker than wild type after 48 hours on glucose but that is probably due to the reduced age of the mutant. Deletion of individual ETC components could drive mitochondrial mis-function rather than simply preventing respiration so we also blocked mitochondrial biogenesis by removing the regulatory Snf4 subunit of the upstream Snf1 complex (yeast AMPK)(Schuller, 2003). Loss of Snf4 did not substantially impair log phase growth in galactose and we were able to purify a 24 hour fraction (albeit with very low cell number as cells died very young), and *snf4*Δ mutants accumulated more Tom70-GFP than wild-type during 24 hours replicative ageing in galactose (Figure 3D, S3B). A plot of Tom70-GFP against mRNA slope shows that *cox9*Δ *gal80*Δ on galactose approaches wild-type on glucose at 48 hours despite little effect at 24 hours, while *snf4*Δ at 24 hours shows an increase in both parameters compared to wild-type at 24 hours on galactose, though far less than wild type on glucose at 24 hours (Figure 3E).

Taken together, these results show that respiration protects cells against the SEP and against gene expression dysregulation, as well as being required for normal lifespan.

### Respiration can protect a subset of cells during ageing in glucose

Yeast can be forced to respire on glucose by over-expression of the mitochondrial biogenesis transcription factor Hap4, which is known to favour ageing on a longer-lived trajectory with rapid cell division to very old age (Lascaris et al., 2003; Li et al., 2020; Lin et al., 2002). In keeping with this report, median Tom70-GFP was significantly lower at 24 hours in Hap4-overexpressing MEP cells on glucose (Figure 4A), but the difference was smaller and non-significant by 48 hours while dysregulation of gene expression was not noticeably rescued (Figure S4A). However, the Tom70-GFP profile of 48 hour-aged Hap4 over-expressing cells was clearly split into two populations, which resolved into a low Tom70-GFP, high WGA population and a high Tom70-GFP, low WGA population akin to the populations observed in wild type cells by Li *et al* (Figure 4B) (Li et al., 2020).

**Figure 4:**
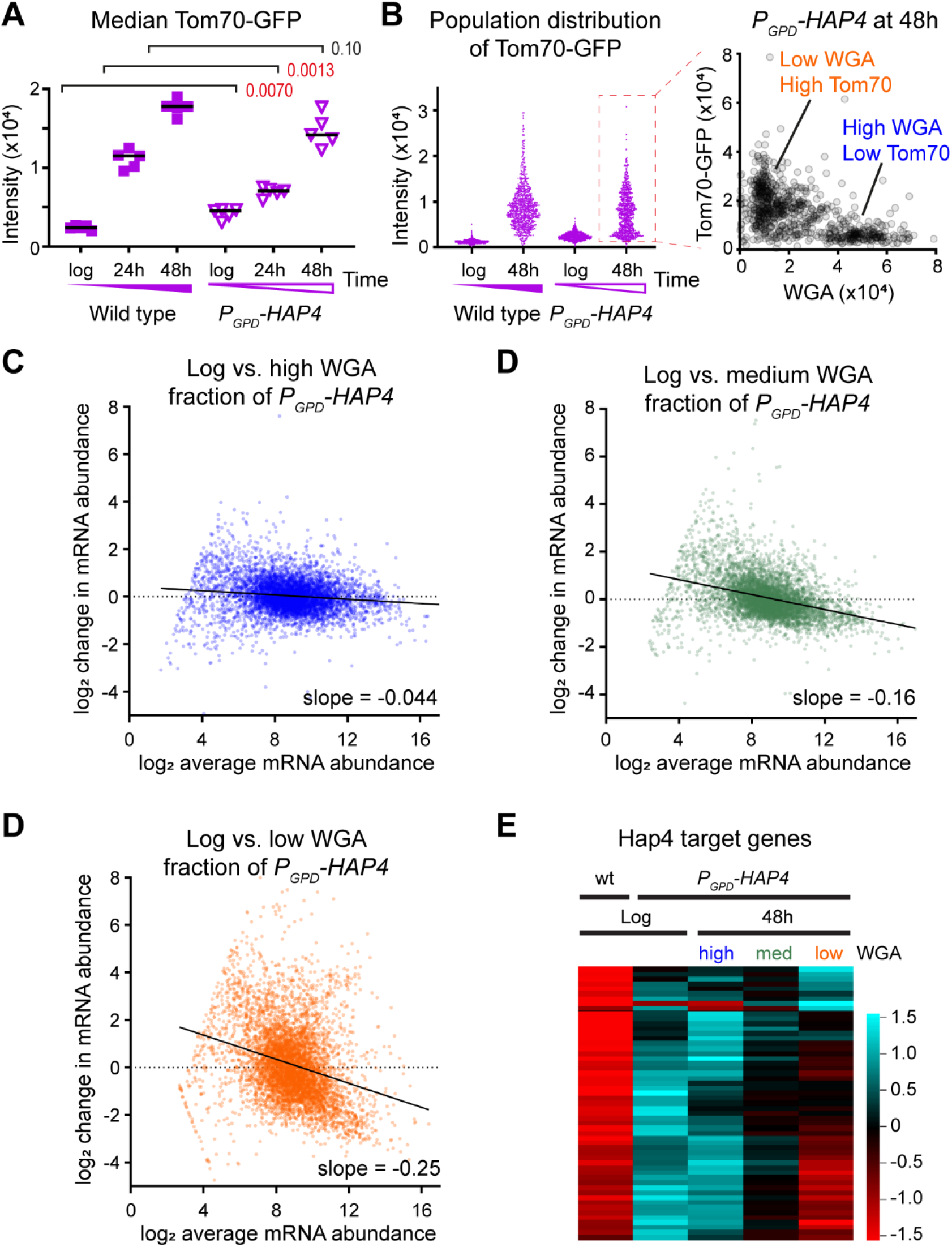
RNA-seq analysis of cells ageing on different trajectories. A: Median Tom70-GFP of wild type and Hap4-overexpressing cells (*P*_*GPD*_*-HAP4*) on glucose, analysed as in Figure 1C, n=5. B: Tom70-GFP population distributions from an individual biological replicate, and plot of Tom70-GFP versus WGA for the *P*_*GPD*_*-HAP4* 48 hour-aged population, highlighting sub-populations demarked by high Tom70-GFP, low WGA (orange) and low Tom70-GFP, high WGA (blue). C: MA plots showing change in mRNA abundance from log phase to the high WGA (left, blue) or low WGA (right, orange) sub-populations of 48 hour-aged *P*_*GPD*_*-HAP4* cells purified by flow cytometry. Cut-off for low expressed genes on which quantification is unreliable at 16 normalised counts, average of 2 biological replicates. D: Heatmap for Hap4-regulated genes (defined as those expressed >3-fold higher in *P*_*GPD*_*-HAP4* compared to wild type at log phase in glucose), for high and low WGA sub-populations of 48 hour aged *P*_*GPD*_*-HAP4* cells compared to log phase wild type and *P*_*GPD-*_*HAP4* controls. Average of 2 biological replicates. E: MA plots showing change in mRNA abundance from log phase to the high WGA (left, blue) or low WGA (right, orange) sub-populations of 48 hour-aged wild-type cells purified by flow cytometry. Cut-off for low expressed genes on which quantification is unreliable at 16 normalised counts. n=1. F: Heatmap for Hap4-regulated genes (defined as those expressed >3-fold higher in *P*_*GPD*_*-HAP4* compared to wild type at log phase in glucose), for high and low WGA sub-populations of 48 hour aged wild-type cells compared to log phase wild type and *P*_*GPD-*_*HAP4* controls. n=1.

In fact, flow cytometry using a BD Influx system detected 3 well-defined fractions in 48 hour-aged Hap4 overexpressing populations: high, medium and low WGA, which we purified and analysed by RNA-seq (Figure S4B). Compared to log phase, the high WGA fraction maintained a youthful gene expression profile with little dysregulation of gene expression, the medium WGA fraction showed an mRNA abundance profile akin to 48 hour aged wild-type, but the low WGA fraction displayed the strongest dysregulation of mRNA abundance we have yet observed (Figure 4C-E). Even after quantile normalisation the low and high WGA profiles are too different for meaningful genome-wide comparisons to identify individual differentially expressed genes (Figure S4C), so we asked whether Hap4 over-expression or that of downstream factors is maintained with age. *HAP4* mRNA itself decreased with age equivalently between the high and low WGA fractions (Figure S4D), but the abundance of many mRNAs that are induced >4-fold by Hap4 over-expression returns to wild-type levels in the low WGA fraction, showing that Hap4 transcription factor activity is not maintained in these cells and therefore the protective effects of respiration are lost (Figure 4F).

We performed an equivalent experiment in wild-type cells to determine whether this heterogeneity is simply a product of Hap4 over-expression (Figure S4E). Although we could not identify defined populations, mRNA abundance profiles of low and high WGA fractions were closely akin to those of low and high WGA fractions from Hap4 overexpressing cells (Figure S4F). This means that within a population aged for a specific time, the youngest cells – those which have ceased to divide rapidly and can be considered senescent – show aberrant mRNA abundance patterns, and therefore gene expression dysregulation must be associated with the SEP rather than ageing itself.

Together, these experiments show that the Hap4 overexpression can increase the fraction of cells that do not undergo SEP onset on glucose, and these cells are marked by youthful gene expression profiles. Combined with the preceding data this means that respiration during ageing suppresses the SEP and age-linked dysregulation of gene expression.

### Accumulation of a chromosomal fragment is tightly linked to the SEP

Transcriptome data includes expression information for rDNA-encoded ncRNAs, which have a major role in rDNA stability and reflect the activity of silencing factors such as Sir2 (Houseley et al., 2007; Kobayashi and Ganley, 2005; Li et al., 2006; Santangelo et al., 1988; Smith and Boeke, 1997). Gene expression patterns in this region are complex so we defined a set of four probes covering each strand of each rDNA intergenic spacer region for quantitation (Figure 5A). These measure overall rDNA ncRNA expression, though the set of ncRNA species detected was very similar between aged samples (Figure S5A). During ageing in glucose, rDNA ncRNA expression rose to a maximum at 24 hours before declining slightly at 48 hours, whilst on galactose these species were much lower at 24 hours but reached the same level as glucose-aged cells by 48 hours (Figure 5B). The ncRNA IGS-1F becomes the most abundant polyadenylated RNA in the cells by 48 hours in glucose and galactose (Figure S5B), and this high expression reflects an equivalent accumulation of ERCs (Figure 5B), which are the primary source of rDNA ncRNA transcripts in aged cells (Pal et al., 2018).

**Figure 5:**
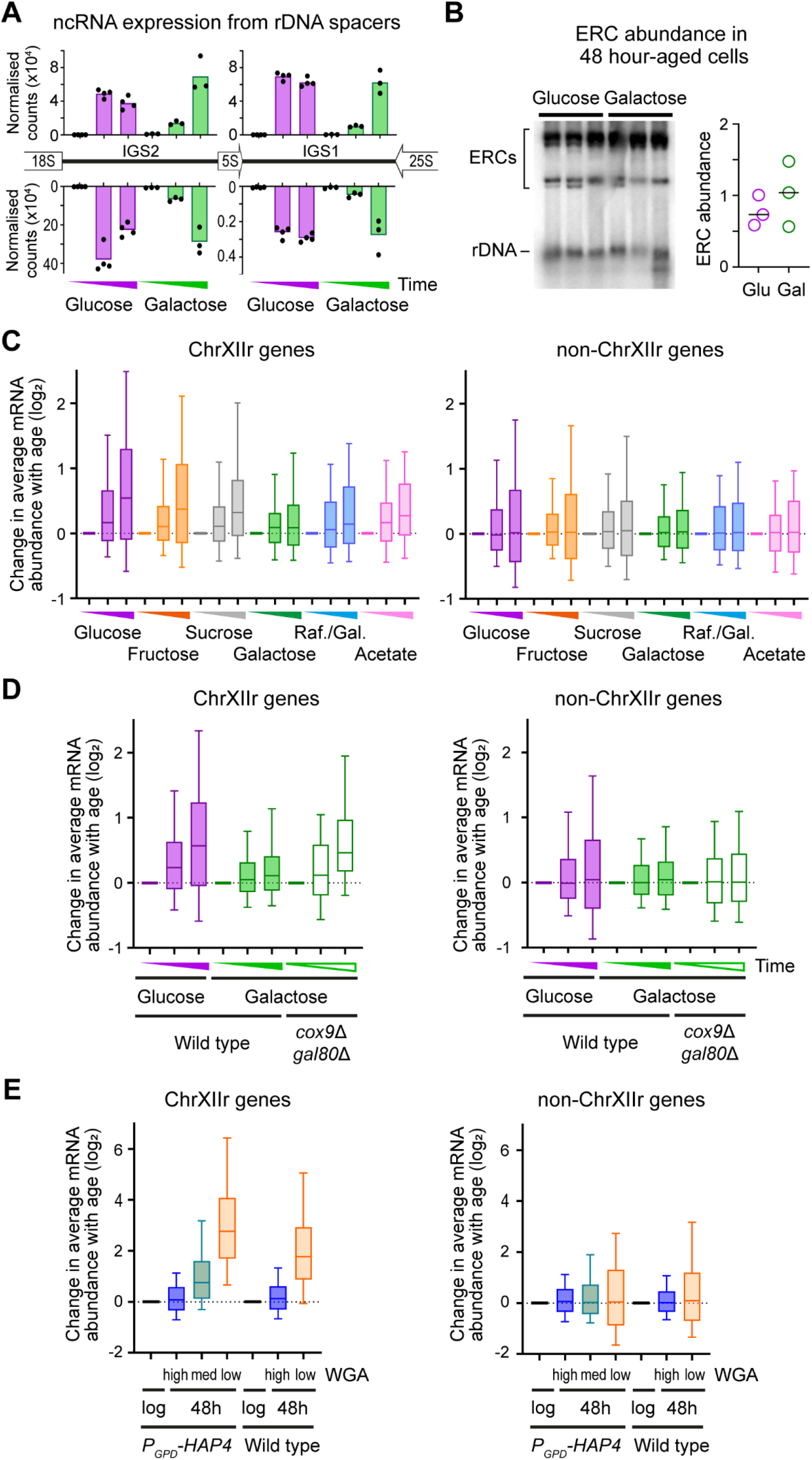
Accumulation of the ChrXIIr fragment is tightly associated with the SEP. A: Quantification of ncRNA transcribed from each strand of the rDNA intergenic spacer regions during ageing on glucose and galactose. Normalised read counts are given for forward and reverse strand transcripts of intergenic spacers IGS1 and IGS2. n=4 for glucose, 3 for galactose. B: Southern blot analysis of ERCs and chromosomal rDNA in 3 biological replicates each of wildtype cells aged for 48 hours on glucose and galactose. Quantification is of the ratio between ERCs and chromosomal DNA. C: Log_2_ change in expression with age of all genes on ChrXIIr [XII:498053-1078177] (left) and all genes except those on chromosome XII and the mitochondrial DNA (right). Boxes show interquartile range with median, whiskers show 10-90th percentiles. n=2 for glucose, fructose and galactose, n=1 for sucrose, raffinose / galactose mix and acetate. D: Change in expression with age of genes on ChrXIIr and controls for wild type on glucose and galactose, and *cox9*Δ *gal80*Δ on galactose. n=2, analysis as in C. E: Change in expression for genes on ChrXIIr between purified fractions of *P*_*GPD*_*-HAP4* and wild type cells aged 48 hours on glucose (Figure 4) relative to log phase controls. n=2 for *P*_*GPD*_*-HAP4*, n=1 for wild type, analysis as in C but note change in scale on y-axis as differences are much greater in these samples.

In the accompanying manuscript Zylstra *et al*. we describe the close association between accumulation of the ChrXIIr fragment (a region of chromosome XII from the rDNA to the centromere-distal telomere) and entry to senescence marked by high Tom70-GFP fluorescence (Hu et al., 2014)(Zylstra *et al*. bioRxiv doi: https://doi.org/10.1101/2022.07.14.500009). This association emerged from analysis of gene expression, and ChrXIIr accumulation can be readily detected by an average increase in mRNA abundance for genes across this region with age. Expression of genes on ChrXIIr increases on average with age in glucose, and (more slowly) fructose and sucrose, while changes in expression of genes in other genomic regions are unbiased (Figure 5C). In contrast, ChrXIIr accumulation was minimal for cells aged on galactose or a raffinose / galactose mixture, while acetate again showed an intermediate phenotype (Figure 5C). These observations reinforce the connection between ChrXIIr accumulation and the SEP, and also mark an association between ChrXIIr and gene expression dysregulation. Furthermore, *cox9*Δ *gal80*Δ mutants ageing on galactose accumulate almost the same level of ChrXIIr by 48 hours on galactose as wild type cells ageing on glucose, in keeping with their increased Tom70-GFP and transcriptional dysregulation (Figures 5D).

Importantly, the low and high WGA fractions of 48 hour aged cells over-expressing Hap4 were differentiated by a massive increase and no change in expression of ChrXIIr genes respectively, meaning that even within a population the presence of ChrXIIr is heterogeneous and specific to the sub-population of cells that have passed the SEP (Figure 5E). In contrast, IGS transcript levels indicative of ERC abundance were equivalent between WGA fractions, and non-chromosome XII regions were largely unaffected (Figure 5E, Figure S5C). Finally, the same division in expression change of ChrXIIr genes was observed between high and low WGA wild-type cell fractions (Figure 5E) meaning that this effect is not caused by over-expression of Hap4 or downstream targets.

These results show that unlike ERCs, accumulation of ChrXIIr is tightly associated with the SEP across a wide range of conditions and mutants, and that dysregulation of gene expression accompanies ChrXIIr accumulation and the SEP.

### Early life events define ageing trajectory on glucose and galactose

Li *et al* demonstrated that the decision between ageing trajectory for cells in glucose is made early in life (Li et al., 2020), and we asked if this principal applies more widely to the time at which diet affects ageing trajectory. We therefore performed media shift experiments, with cells aged for 24 hours in glucose or galactose (at which point viability is still >80%, Figure 1D), then harvested and resuspended either in media containing the same or the reciprocal sugar (Figure 6A).

**Figure 6:**
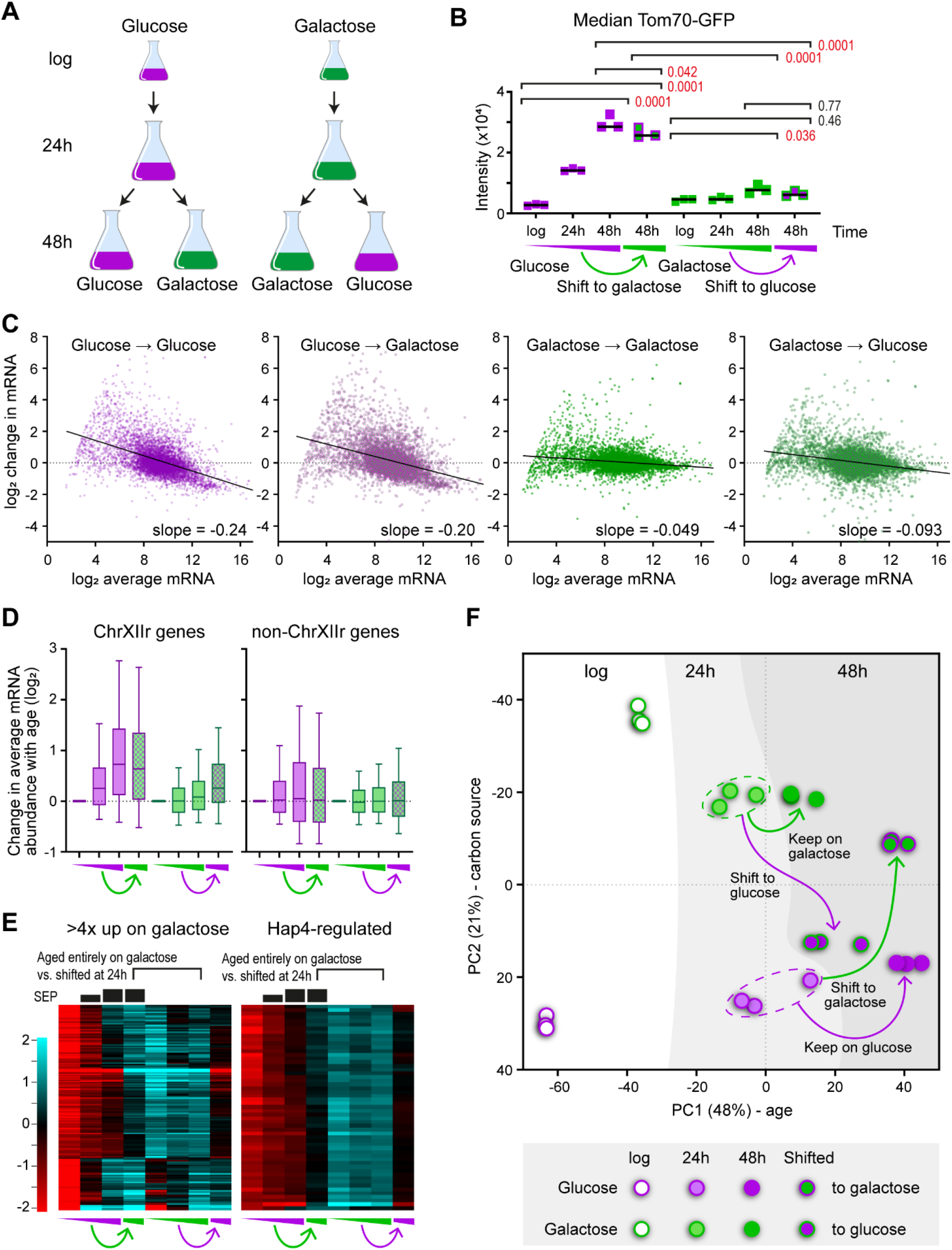
Media shifts determine the plasticity of ageing phenotypes. A: Schematic of experimental set up. Cells are harvested at 24 hours then transferred to either the same or the reciprocal media. B: Median Tom70-GFP intensities for all populations, analysed as in Figure 1C. p values from 1 way ANOVA with post-hoc Tukey test, n=3. C: MA plots for all four 48 hour aged fractions compared to the relevant log phase controls, performed as in Figure 2A, average of 3 biological replicates. D: Change in expression with age for genes on ChrXIIr and control set as Figure 5C, average of 3 biological replicates. E: Heatmap across all samples of genes either induced >4x on galactose relative to glucose at log phase (left) or of the Hap4-regulated gene set from Figure 4D (right). The extent of SEP entry is indicated diagrammatically above the plots based on Tom70-GFP intensities from B. Average of 3 biological replicates. F: PCA plot showing changes in mRNA expression pattern, as Figure 2C.

The SEP, as marked by Tom70 intensity, was dependent on early life diet (Figure 6B, Figure S6A): if the first 24 hours were spent on glucose, the Tom70-GFP signal observed at 48 hours was very high and ageing on galactose for the second 24 hours had a significant but not substantial effect; in contrast if the first 24 hours were spent on galactose, ageing on glucose for the second 24 hours did not cause an increase in Tom70-GFP signal. Unsurprisingly given the previous results, gene expression dysregulation was strong in populations that spent the first 24 hours on glucose and much lower for those that spent the first 24 hours in galactose, irrespective of diet for the second 24 hours (Figure 6C). Furthermore, ChrXIIr accumulation was detected only if the first 24 hours were spent in glucose (Figure 6D), whereas rDNA ncRNA expression was similar between all 48 hour aged samples, though lower at 24 hours in galactose than in glucose (Figure S6B).

It has been reported that ageing cells lose the capacity to properly induce genes, which could explain the importance of the first 24 hours (Neurohr et al., 2018). However, the set of genes that are expressed >4-fold higher in galactose than glucose at log, and are presumably critical for metabolic state, are mostly induced to the same level at 48 hours irrespective of whether the first 24 hours of ageing were spent on glucose or galactose (Figure 6E, left). We see no difference in steady state levels of the core galactose regulon of *GAL1, GAL10, GAL7, GAL2* after media shift at 24 hours (Figure S6C), though induction may of course be slower, but Hap4 target gene expression was lower in cells shifted from glucose to galactose at 24 hours compared to cells that were aged entirely on galactose (Figure 6E, right). Nonetheless, Hap4 target gene expression is not the defining factor for senescence as expression of these genes is if anything slightly higher in glucose-to-galactose shifted cells that exhibit high Tom70-GFP, compared to galactose-to-glucose shifted cells that exhibit low Tom70-GFP (Figure 6E, right).

Globally, a PCA of the transcriptome data divides the influence of diet in the first and second 24 hours (Figure 6F). Distance travelled on PC1 is defined by the first 24 hours with cells that start on galactose travelling a shorter distance than those which start on glucose irrespective of diet for the second 24 hours, so the glucose→glucose and glucose→galactose samples are coincident on PC1, as are the galactose→galactose and galactose→glucose samples. However, PC2 is divided based on diet at point of harvesting, so the glucose log and 24 hours are coincident on PC2 with the glucose→glucose and galactose→glucose samples, ie: it is the diet for the second 24 hours that affects PC2. This separation also occurs for samples on galactose at point of harvest; these are all clearly segregated from glucose on PC2 although spread over a larger range of PC2 values. Therefore, highly aged cells largely maintain responsiveness of gene expression to dietary change late in life, but gene expression dysregulation is defined by diet early in life.

Overall, our study shows that diet in early life controls the appearance of age-linked phenotypic changes, even without dietary restriction. This arises through respiration suppressing onset of the SEP, with successful induction of respiration redirecting cells into a pathway that minimises key hallmarks of ageing even on glucose.

## Discussion

The ability to redirect cells from an unhealthy to a healthy ageing trajectory is the primary aim of research into healthy ageing. Here we demonstrate that substitution of galactose for glucose suppresses cell intrinsic processes that lead to a spectrum of seemingly pathological changes during yeast replicative ageing. This does not extend lifespan, showing that the mechanisms which limit lifespan are separable from those that cause senescence.

### Diet-induced transitions between ageing trajectories

It is often assumed that ageing pathology and eventual death are the result of a progressive accumulation of unavoidable damage. However, in yeast we find that senescence is not obligatory and that the extent of replicative lifespan is not dependent on damage accumulation, at least not any form of damage that we have assayed. Instead, our findings are consistent with models in which ageing can follow different trajectories, only some of which are associated with pathology. Other trajectories do not result in immortality, but the cause of death is not the accumulation of senescence marks and remains unknown. Importantly, such trajectories end with loss of viability without an extended period of senescence, and can therefore be considered as healthy ageing.

Ageing trajectory is defined early in life for yeast and is enforced by a mechanism resilient to changes in gene expression and metabolic state, since media shifts at mid-life do not curtail the onset of the SEP, gene expression dysregulation or ChrXIIr accumulation. Extrachromosomal DNA retained in the mother cell such as ERCs or ChrXIIr offer an insufficient explanation: ERCs do not correlate to any of these phenotypes, and ChrXIIr continues to accumulate after the media shift so is likely a consequence rather than a cause of the trajectory. We doubt that any epigenetic mark at the rDNA or elsewhere would be sufficiently stable or asymmetrically segregated to maintain this trajectory. Loss of vacuolar acidity has been proposed to drive ageing in yeast and could force cells permanently into the pathological trajectory, as the vacuole regulates iron availability and a lack of iron causes numerous cellular defects (Chen et al., 2020; Hughes and Gottschling, 2012; Ramos-Alonso et al., 2020; Veatch et al., 2009). Iron availability is low in the short-lived trajectory, iron importer subunits are upregulated both in single cell RNA-seq of slow dividing ageing cells and in our RNA-seq analysis of low WGA cells, and we observe that over-expression of the iron-dependent Hap4 transcription factor cannot maintain expression of target genes in cells on the pathological ageing trajectory (Li et al., 2020; Zhang et al., 2020). These observations show that iron deficiency is a feature of the short-lived, high SEP trajectory, but the reason why respiration is so effective in removing cells from this trajectory, and the reason why only a fraction of wild-type cells on glucose follow this trajectory, remain unclear.

Unfortunately, even in yeast the relationship between diet and ageing is metastable; results that are highly reproducible within one experimental system are not necessarily reproduced in others. For example, 2%→0.5% caloric restriction robustly extends lifespan in microdissection experiments but does not extend lifespan in microfluidic studies (Huberts et al., 2014; Jo et al., 2015; Lee et al., 2012), and similarly galactose improves ageing cell fitness when assayed by microdissection but not by microfluidics (Aspert and Charvin, personal communication)(Frenk et al., 2017). From this viewpoint, it is very helpful that the data for ageing trajectories encompasses microfluidics, microdissection and now batch culture experiments performed on different derivatives of a standard genetic background (S288C) by different groups, and is therefore robust across experimental systems. This suggests that different ageing trajectories are an intrinsic part of the yeast ageing process with environmental and genetic manipulations biasing the process towards different trajectories, and therefore even if the effects of specific environmental changes are not reproducible across all experimental systems, the underlying biology is likely to be consistent.

In trying to understand ageing trajectories, it is worthwhile considering potential cause and effect relationships between the ageing phenotypes we have examined, both here and in Zylstra *et al*. Formation of intense Tom70-GFP foci in G1 cells post SEP is very tightly associated with ChrXIIr across all our datasets, but given that ChrXIIr will persist throughout the cell cycle, it seems most likely that ChrXIIr formation is a proximal cause of the Tom70-GFP phenotype not vice versa. This idea fits very well with the observation of Neurohr *et al*. that the transfer of extrachromosomal DNA to daughter cells transmits the senescent phenotype (Neurohr et al., 2018). In contrast, gene expression dysregulation is unaffected by reduced ChrXIIr accumulation in *spt3*Δ and remains low in *rad52*Δ despite abundant ChrXIIr and early SEP onset (Zylstra *et al*. bioRxiv doi: https://doi.org/10.1101/2022.07.14.500009). This means that gene expression dysregulation lies either upstream or occurs independently of the SEP, but the tight segregation in gene expression dysregulation within populations between SEP and non-SEP cells strongly suggests that gene expression dysregulation is an early mechanistic driver of the SEP. Under such a model, mutations such as *spt3*Δ and *rad52*Δ simply modulate the rapidity with which underlying ageing phenotypes, of which gene expression dysregulation may only be one, manifest in senescence phenotypes.

### The importance of respiration in healthy ageing

The benefits of dietary restriction have been attributed to respiration (Anderson et al., 2003; Lin et al., 2004; Lin et al., 2002; Schleit et al., 2013), and we find that the health benefits of a galactose diet similarly require an active electron transport chain. Respiration is obligatory in higher eukaryotes, but declining expression of mitochondrial protein genes is the most robust feature of ageing transcriptomes across higher eukaryotes and one that is rapidly reversed under dietary restriction (reviewed in (Frenk and Houseley, 2018)).

Although enhanced respiration remains disputed as the mechanism of action for dietary restriction in yeast (Kaeberlein et al., 2004), forced respiration during growth on glucose extends lifespan and promotes more cells to a healthy ageing trajectory (Li et al., 2020; Lin et al., 2002; Patnaik et al., 2022). Our findings reinforce these observations and show that cells pushed into this trajectory by overexpression of Hap4 maintain a youthful gene expression state to very advanced age. Loss of iron-dependent Hap4 activity, which likely results from the loss of iron homeostasis in a fraction of cells during ageing on glucose (Chen et al., 2020; Li et al., 2020; Zhang and Hach, 1999; Zhang et al., 2020), is observed in Hap4 over-expressing cells that do follow the healthy ageing trajectory. However, Hap4 target genes, which include many critical respiration proteins, were not differentially expressed between cells that have shifted carbon source at 24 hours despite dramatically different ageing trajectory. This means that expression of Hap4 target genes is either sufficient but not necessary for cells to enter the long lived, healthy ageing trajectory, or that Hap4 target gene expression is only necessary early in life.

Across this study and the accompanying manuscript, the molecular change most strongly associated with the SEP is ChrXIIr fragment accumulation. In Zylstra *et al*., we propose that ChrXIIr forms as a result of replication for stalling in the rDNA, which would be promoted in the absence of Sir2 (Zylstra *et al*., bioRxiv, doi: https://doi.org/10.1101/2022.07.14.500009). The activity of the histone deacetylase Sir2 is promoted by respiration through the formation of NAD+ and clearance of nicotinamide (Lin et al., 2004), so it is reasonable to relate rDNA recombination events that form ChrXIIr to a lack of respiration. It should be noted that others have attributed the effect of caloric restriction on lifespan to reduced activity of mTOR rather than respiration (Kaeberlein et al., 2004); but this could have a similar outcome as mTOR signalling promotes rDNA recombination through Sir2 and the homologous histone deacetylases Hst3 and Hst4 (Ha and Huh, 2011; Jack et al., 2015; Medvedik et al., 2007). Therefore, multiple pathways affected by diet coincide on the rDNA locus, all of which could alter the production of the ChrXIIr fragment that is so tightly linked to the SEP.

Overall, our study shows that a transition to a constitutively healthy ageing trajectory is possible, at least in yeast. Of course, substituting galactose as the primary caloric input is neither achievable nor useful in humans, but our findings suggest that dietary change without restriction can offer paths to ageing health benefits currently only observed under dietary restriction.

## Materials and Methods

Detailed and updated protocols are available at https://www.babraham.ac.uk/our-research/epigenetics/jon-houseley/protocols

### Yeast culture and labelling

Yeast strains were constructed by standard methods and are listed in Table S1, oligonucleotide sequences are given in Table S2. Plasmid templates were pFA6a-GFP-KanMX4, pYM-N14, pAW8-mCherry, pFA6a-HIS3 and pFA6a-URA3 (Houseley and Tollervey, 2011; Janke et al., 2004; Longtine et al., 1998; Watson et al., 2008). All cells were grown in YPx media (2% peptone, 1% yeast extract, 2% sugar) at 30°C with shaking at 200 rpm. Media components were purchased from Formedium and media was sterilised by filtration. MEP experiments were performed as described with modifications (Cruz et al., 2018): cells were inoculated in 4 ml YPx (where x is the sugar) and grown for 6-8 hours then diluted in 25 ml YPx and grown for 16-18 hours to 0.2-0.6×10^7^ cells/ml in 50 ml Erlenmeyer flasks. 0.125×10^7^ cells per aged culture were harvested by centrifugation (15 s at 13,000 g), washed twice with 125 µl PBS and re-suspended in 125 µl of PBS containing ∼3 mg/ml Biotin-NHS (Pierce 10538723). Cells were incubated for 30 min on a wheel at room temperature, washed once with 125 µl PBS and re-suspended in 125 µl YPx then inoculated in 125 ml YPx at 1×10^4^ cells/ml in a 250 ml Erlenmeyer flask (FisherBrand FB33132) sealed with Parafilm. 1 µM β-estradiol (from stock of 1 mM Sigma E2758 in ethanol) was added after 2 h (glucose, fructose), 3 h (galactose, sucrose) or 4 h (potassium acetate). An additional 0.125×10^7^ cells were harvested from the log phase culture while biotin labelling reactions were incubating at room temperature. Cells were harvested by centrifugation for 1 min, 4600 rpm, immediately fixed by resuspension in 70% ethanol and stored at -80°C. To minimise fluorophore bleaching in culture, the window of the incubator was covered with aluminium foil, lights on the laminar flow hood were not used during labelling and tubes were covered with aluminium foil during biotin incubation. Lifespan assays and competition assays in the MEP background were performed as previously described (Cruz et al., 2018; Frenk et al., 2017).

### Cell purification

Percoll gradients (1-2 per sample depending on harvest density) were formed by vortexing 1 ml Percoll (Sigma P1644) with 110 µl 10x PBS in 2 ml tubes and centrifuging 15 min at 15,000 g, 4 °C. Ethanol fixed cells were defrosted and washed once with 1 volume of cold PBSE (PBS + 2 mM EDTA) before resuspension in ∼100 µl cold PBSE per gradient and layering on top of the pre-formed gradients. Gradients were centrifuged for 4 min at 2,000 g, 4 °C, then the upper phase and brown layer of cell debris removed and discarded. 1 ml PBSE was added, mixed by inversion and centrifuged 1 min at 2,000 g, 4 °C to pellet the cells, which were then re-suspended in 1 ml PBSE per time point (re-uniting samples where split across two gradients). 25 µl Streptavidin microbeads (Miltenyi Biotech 1010007) were added and cells incubated for 5 min on a wheel at room temperature. Meanwhile, 1 LS column per sample (Miltenyi Biotech 1050236) was loaded on a QuadroMACS magnet and equilibrated with cold PBSE in 4 °C room. Cells were loaded on columns and allowed to flow through under gravity, washed with 1 ml cold PBSE and eluted with 1 ml PBSE using plunger. Cells were re-loaded on the same columns after re-equilibration with ∼500 µl PBSE, washed and re-eluted, and this process repeated for a total of three successive purifications. After addition of Triton X-100 to 0.01% to aid pelleting, cells were split into 2 fractions in 1.5 ml tubes, pelleted 30 s at 20,000 g, 4 °C, frozen on N2 and stored at -70 °C.

### RNA extraction

Cells were re-suspended in 50 µl Lysis/Binding Buffer (from mirVANA kit, Life Technologies AM1560), and 50 µl 0.5 µm zirconium beads (Thistle Scientific 11079105Z) added. Cells were lysed with 5 cycles of 30 s 6500 ms^-1^ / 30 s on ice in an MP Fastprep bead beater or for 3 min at power 12 in a Bullet Blender (ThermoFisher) in cold room, then 250 µl Lysis/Binding buffer added followed by 15 µl miRNA Homogenate Additive and cells were briefly vortexed before incubating for 10 minutes on ice. 300 µl acid phenol : chloroform was added, vortexed and centrifuged 5 min at 13,000 g, room temperature before extraction of the upper phase. 400 µl room temperature ethanol and 2 µl glycogen (Sigma G1767) were added and mixture incubated for 1 hour at -30 °C before centrifugation for 15 minutes at 20,000 g, 4 °C. The pellet was washed with cold 70% ethanol and re-suspended in 10 µl water. 1 µl RNA was glyoxylated and analysed on a BPTE mini-gel, and RNA was quantified using a PicoGreen RNA kit (Life Technologies R11490) or Qubit® RNA HS Assay Kit.

150 ng RNA was used to prepare libraries using the NEBNext Ultra II Directional mRNAseq kit with poly(A)+ purification module (NEB E7760, E7490) as described with modifications: Reagent volumes for elution from poly(T) beads, reverse transcription, second strand synthesis, tailing and adaptor ligation were reduced by 50%; libraries were amplified for 13 cycles using 2 µl each primer per 50 µl reaction before two rounds of AMPure bead purification at 0.9x and elution in 11 µl 0.1x TE prior to quality control using a Bioanalyzer HS DNA ChIP (Agilent) and quantification using a KAPA Library Quantification Kit (Roche).

### Sequencing and bioinformatics

Libraries were sequenced by the Babraham Institute Sequencing Facility using a NextSeq 500 instrument on 75 bp single end mode. After adapter and quality trimming using Trim Galore (v0.6.6), RNA-seq data was mapped to yeast genome R64-1-1 using HISAT2 v2.1.0 (Kim et al., 2019) by the Babraham Institute Bioinformatics Facility. Mapped data was imported into SeqMonk v1.47.0 (https://www.bioinformatics.babraham.ac.uk/projects/seqmonk/) and quantified for log_2_ total reads mapping to the antisense strand of annotated open reading frames (opposite strand specific libraries), excluding the mtDNA and the rDNA locus, but with probes included to each strand of the rDNA intergenic spacer regions. Read counts were adjusted by Size Factor normalisation for the full set of quantified probes (Anders and Huber, 2010). Where indicated, a quantile normalisation was additionally applied using the ‘Match Distribution’ function in Seqmonk.

MA plots were generated in GraphPad Prism (v9.2.0) comparing mean and difference for each gene between two conditions. Probes with very low numbers of reads post normalisation in the control condition were filtered (cut-offs given in Figure legends) as change between datasets cannot be accurately quantified in this case. Slope values were determined from the same filtered datasets using the lm(difference∼mean) function in R. PCA plots, hierarchical clustering plots and heatmaps were calculated within SeqMonk, DESeq2 analysis was performed after Quantile normalisation using the Match Distribution function within SeqMonk. GO analysis of individual clusters performed using GOrilla (http://cbl-gorilla.cs.technion.ac.il/) (Eden et al., 2009). Quoted p-values for GO analysis are FDR-corrected according to the Benjamini and Hochberg method (q-values from the GOrilla output), for brevity only the order of magnitude rather than the full q-value is given (Benjamini and Hochberg, 1995). Full GO analyses are provided in Supplementary File S1.

All RNA-seq data has been deposited at GEO under accession number GSE207503. Aged RNA-seq data for many mutants not included in this manuscript is also deposited in this accession.

### Flow cytometry

Cell pellets were re-suspended in 240 µl PBS and 9 μl 10% triton X-100 containing 0.3 µl of 1 mg/ml Streptavidin conjugated with Alexa Fluor® 647 (Life technologies) and 0.6 μl of 1 mg/ml Wheat Germ Agglutinin (WGA) conjugated with CF®405S (Biotium). Cells were stained for 10 min at RT on a rotating mixer while covered with aluminium foil, washed once with 300 μl PBS containing 0.01% Triton X-100, re-suspended in 30 μl PBS and immediately subject to flow cytometry analysis. Flow cytometry analysis was conducted using an Amnis® ImageStream® X Mk II with the following laser power settings: 405=25mW, 488=180mW, 561=185mW, 642=5mW, SSC=0.3mW.

Cell populations were gated for single cells based on Area and Aspect Ratio (>0.8) values and in-focus cells were gated based on a Gradient RMS value (>50). Further gating of streptavidin positive (AF647) cells was also applied, all in a hierarchical manner and 1000 events acquired. Before data analysis, compensation was applied according to single-colour controls and a manual compensation matrix creation. Total fluorescence intensity values of different parameters were extracted using the Intensity feature of the IDEAS® software, with Adaptive Erode modified mask coverage. In the analysis, only positive values of fluorescence were included (i.e. where cells were truly positive for the marker) and median values of populations were determined with Graphpad Prism (v9.2.0). Cell sorting was conducted using a BD Influx™ system. Cell populations were gated for single cells based on forward and side scatter parameters and aged cells based on streptavidin stain. Distinct populations were sorted based on GFP and WGA-AF405 intensities.

### Statistical analysis

All statistical analysis was performed in GraphPad Prism (v9.2.0).

## Acknowledgements

We would like to thank Rachael Walker, Attila Bebes and Aleksandra Lazowska-Addyof the Babraham Institute Flow Facility for their assistance with imaging flow cytometry and sorting of yeast cells, Paula Kokko-Gonzales, Nicole Forrester and Amelia Edwards of the Babraham Institute Next Generation Sequencing facility for sequencing samples and QC assistance, and Simon Andrews, Anne Segonds-Pichon and Felix Krueger of the Babraham Institute Bioinformatics Facility for help with interpreting sequencing data, statistical analysis and sample processing. We thank Grazia Pizza for southern blot analysis, Gilles Charvin and Théo Aspert for helpful discussions, Andre Zylstra and Della David for critical reading, and Patrick Lindstrom and Dan Gottschling for provision of the MEP system.

---

**Figure S1:**
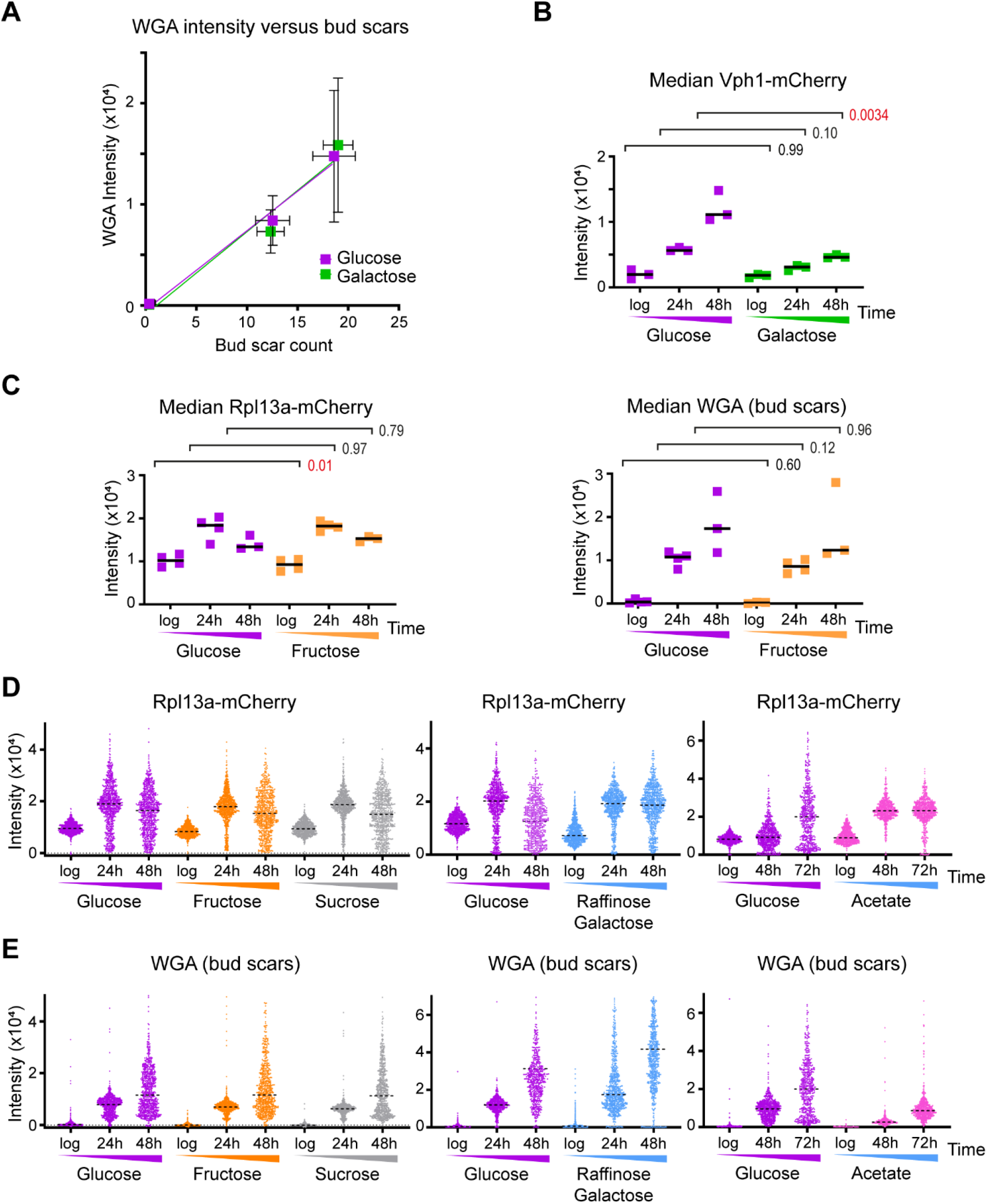
Supplement to progression of age-linked phenotypes on different diets. A: WGA intensities versus manual bud scar counts across ageing on glucose and galactose. B: Median Vph1-mCherry intensities at different ages on glucose and galactose, as Figure 1C, n=3. C: Rpl13a-mCherry and WGA data for samples in Figure 1F. D: Rpl13a-mCherry data for samples in Figure 1G. E: WGA data for samples in Figure 1G.

**Figure S2:**
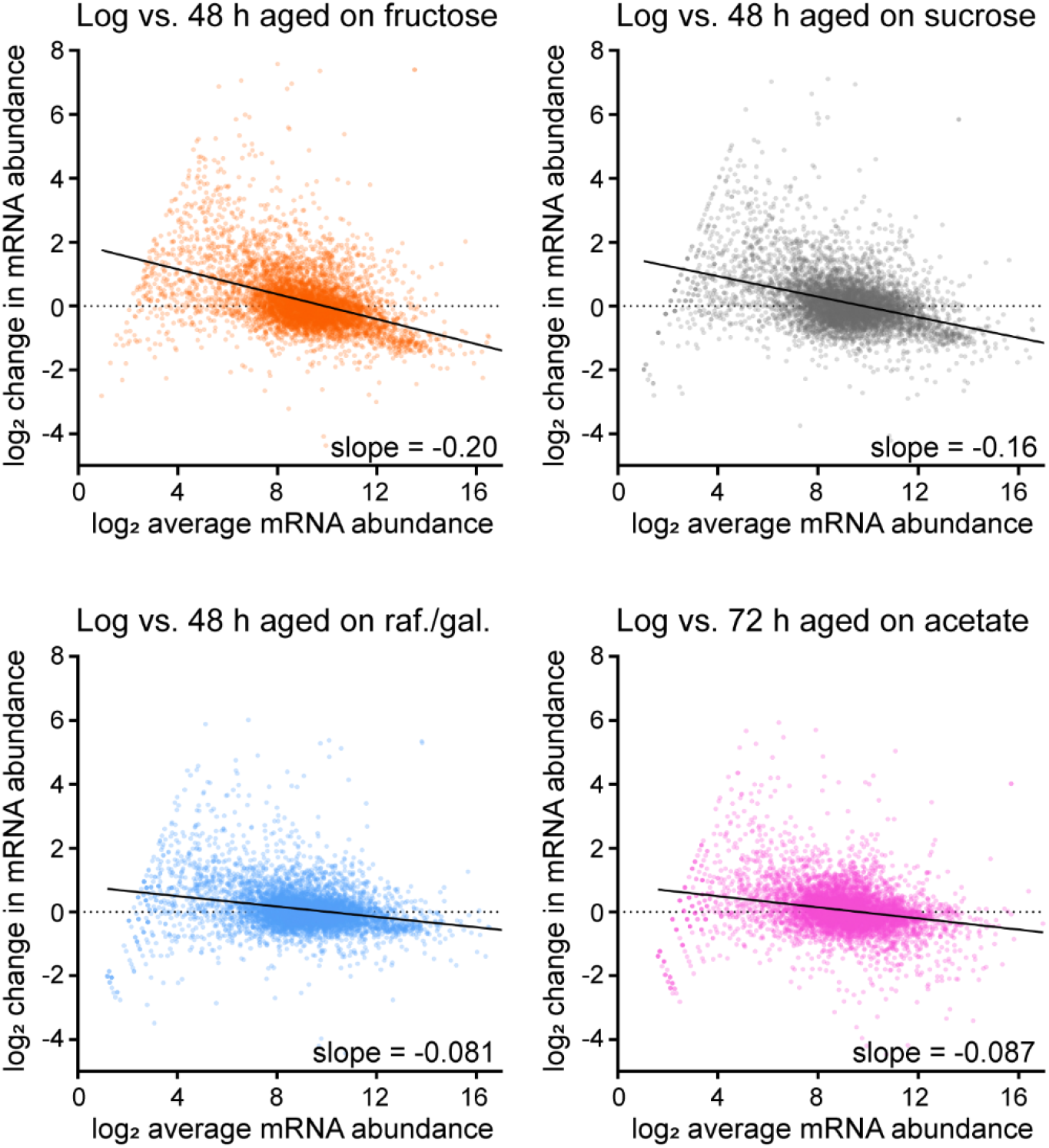
Supplement to effects of diet on gene expression change during ageing. MA plots for gene expression change between log phase and 48 hours ageing for wild type cells on fructose, sucrose or a 2% raffinose 0.5% galactose mixture, and equivalent plot for cells aged for 72 hours on 2% potassium acetate.

**Figure S3:**
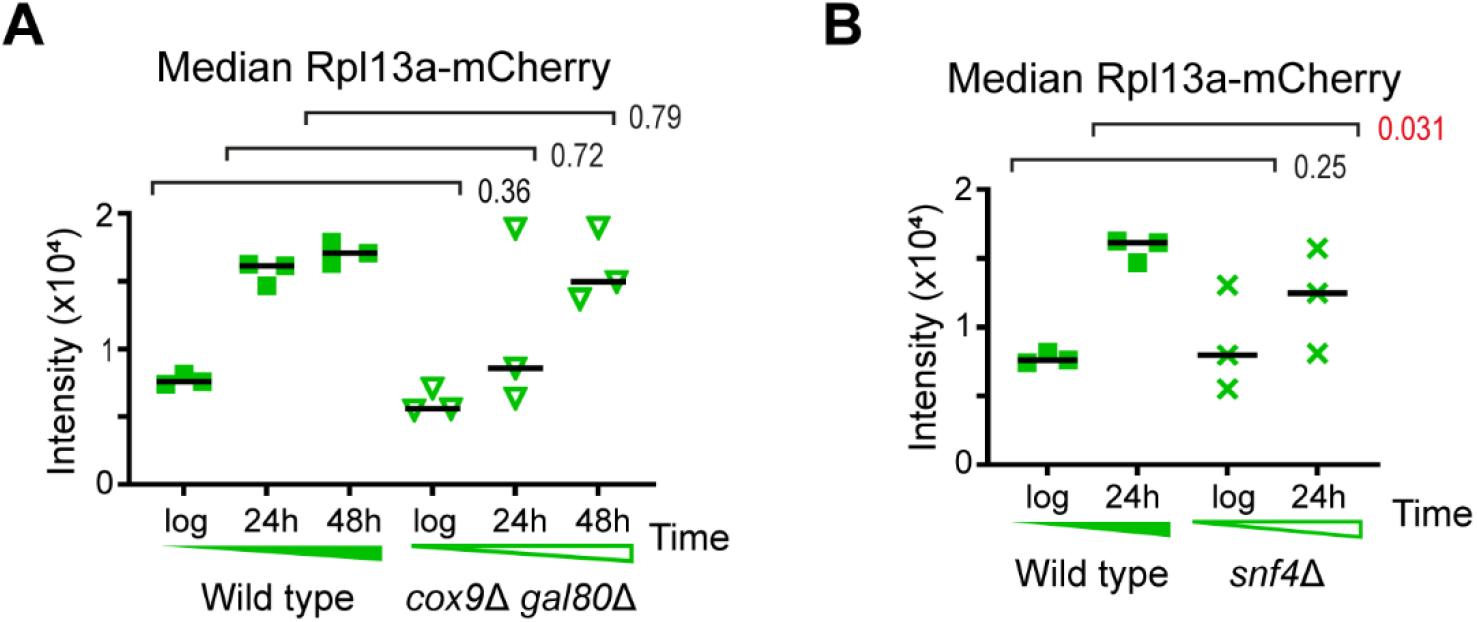
Supplement to respiration is required to avoid the SEP and gene expression dysregulation on galactose. A: Rpl13a-mCherry data for samples in Figure 3B. B: Rpl13a-mCherry data for samples in Figure 3D.

**Figure S4:**
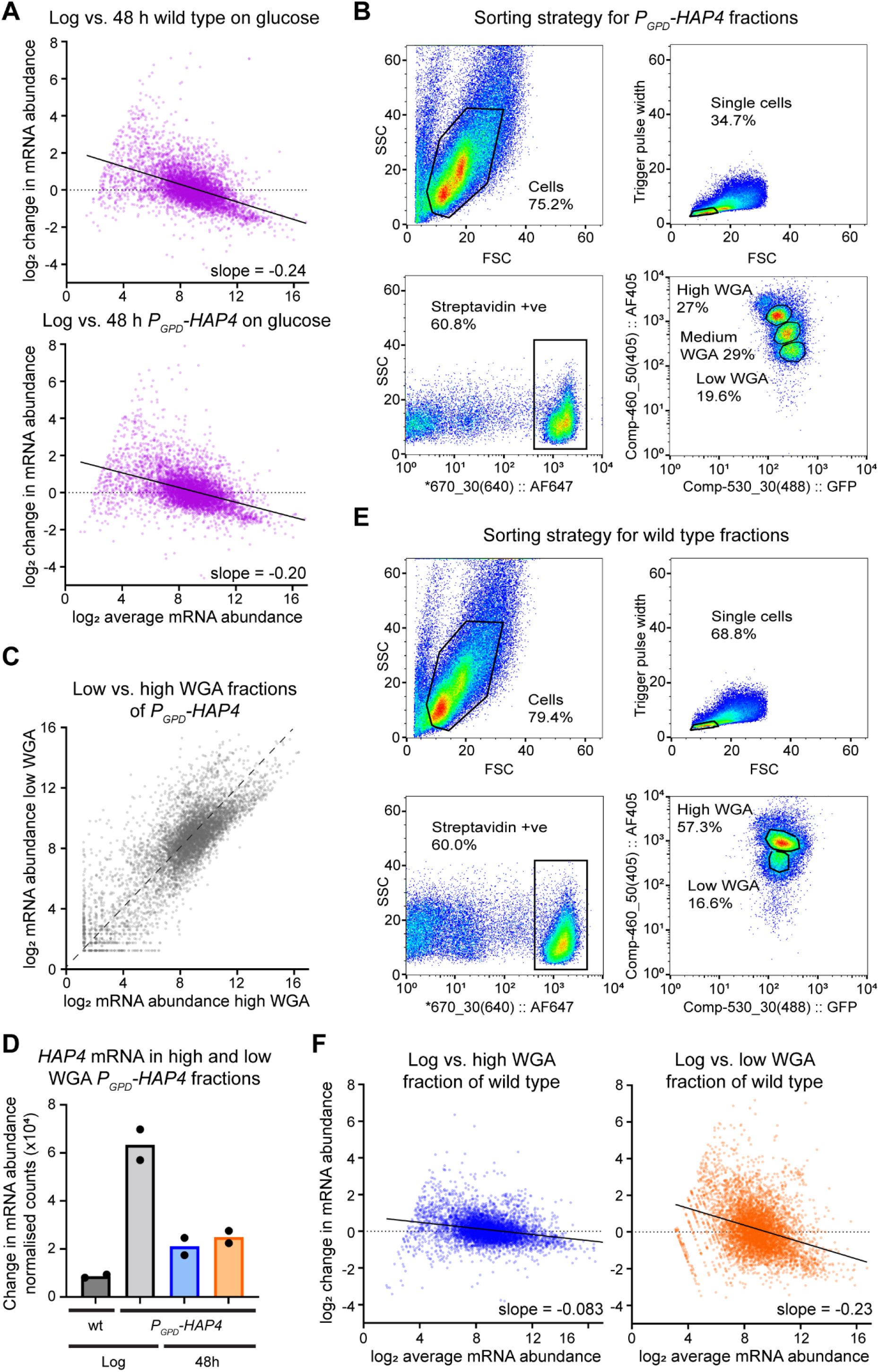
Supplement to RNA-seq analysis of cells ageing on different trajectories. A: MA plots comparing 48 hour-aged on glucose to log phase control samples for wild type (left) and *P*_*GPD*_*-HAP4* (right). Average of 2 biological replicates. B: Flow sorting strategy and gating for *P*_*GPD*_*-HAP4* cells aged for 48 hours on glucose. C: Scatter plot comparing quantile-normalised mRNA abundances in low and high WGA fractions of *P*_*GPD*_*-HAP4* 48 hour-aged cells. D: *HAP4* mRNA abundance in *P*_*GPD*_*-HAP4* 48 hour-aged cells compared to wild-type controls, n=2 biological replicates. E: Flow sorting strategy and gating for wild-type cells aged for 48 hours on glucose. F: Plot of Tom70-GFP versus WGA intensities from a 72 hour-aged wildtype sample on glucose, indicating low WGA, high Tom70-GFP and high WGA, low Tom70-GFP sub-populations.

**Figure S5:**
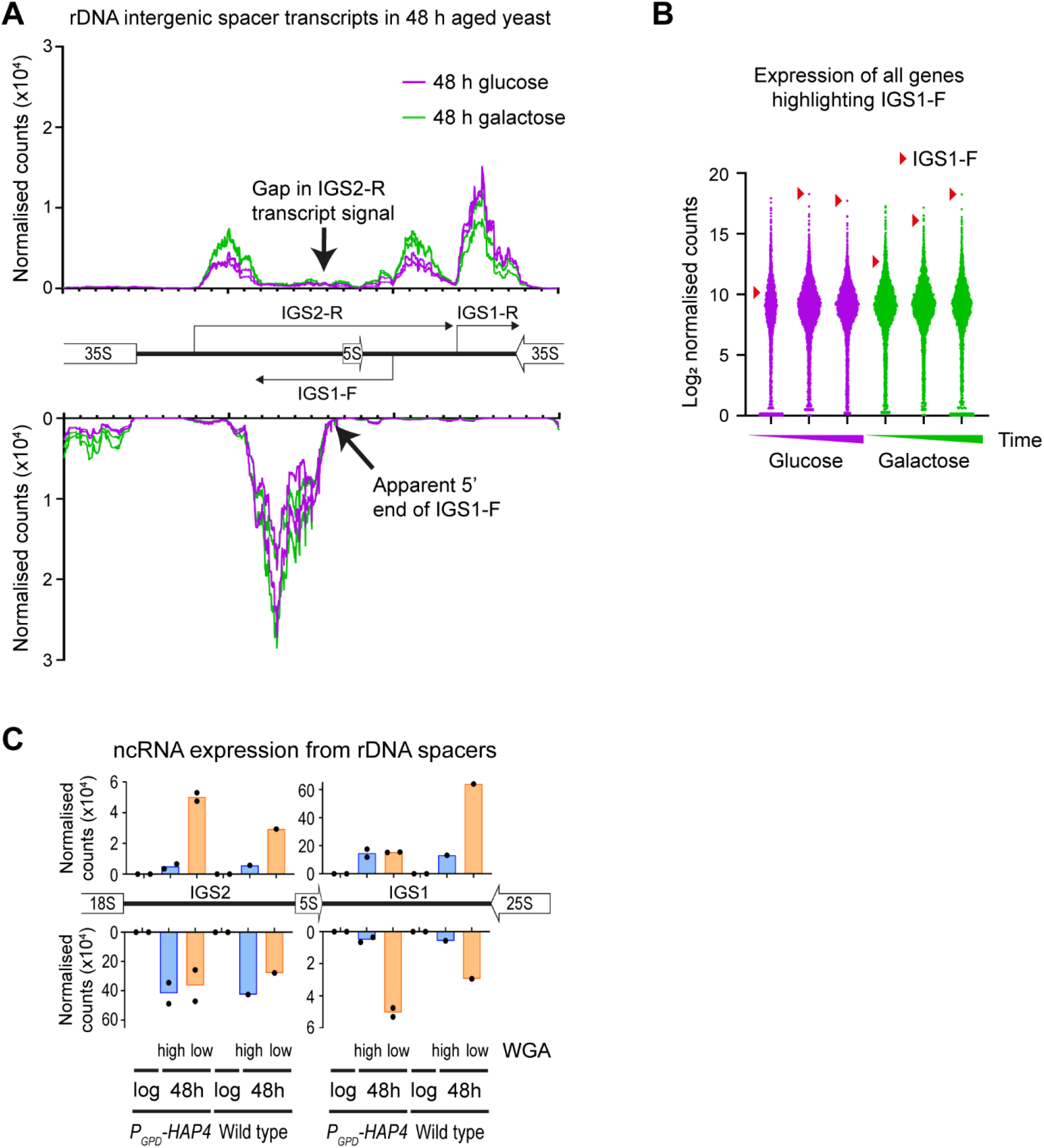
Supplement to accumulation of the ChrXIIr fragment is tightly associated with the SEP. A: RNA read counts for each strand in 10 bp windows across rDNA intergenic spacer regions, for 2 biological replicate samples each on aged for 48 hours on glucose or galactose. Previously described RNA species are indicated (Houseley et al., 2007), along with deviations from expected profiles. B: Log_2_ normalised read counts for all genes including the rDNA intergenic spaces. The individual dot in each sample corresponding to the rDNA intergenic spacer probe containing the IGS1-F ncRNA is indicated by a red arrow. C: rDNA ncRNA abundances in the sorted fractions of *P*_*GFP*_*-HAP4* and wild-type cells from Figure 4.

**Figure S6:**
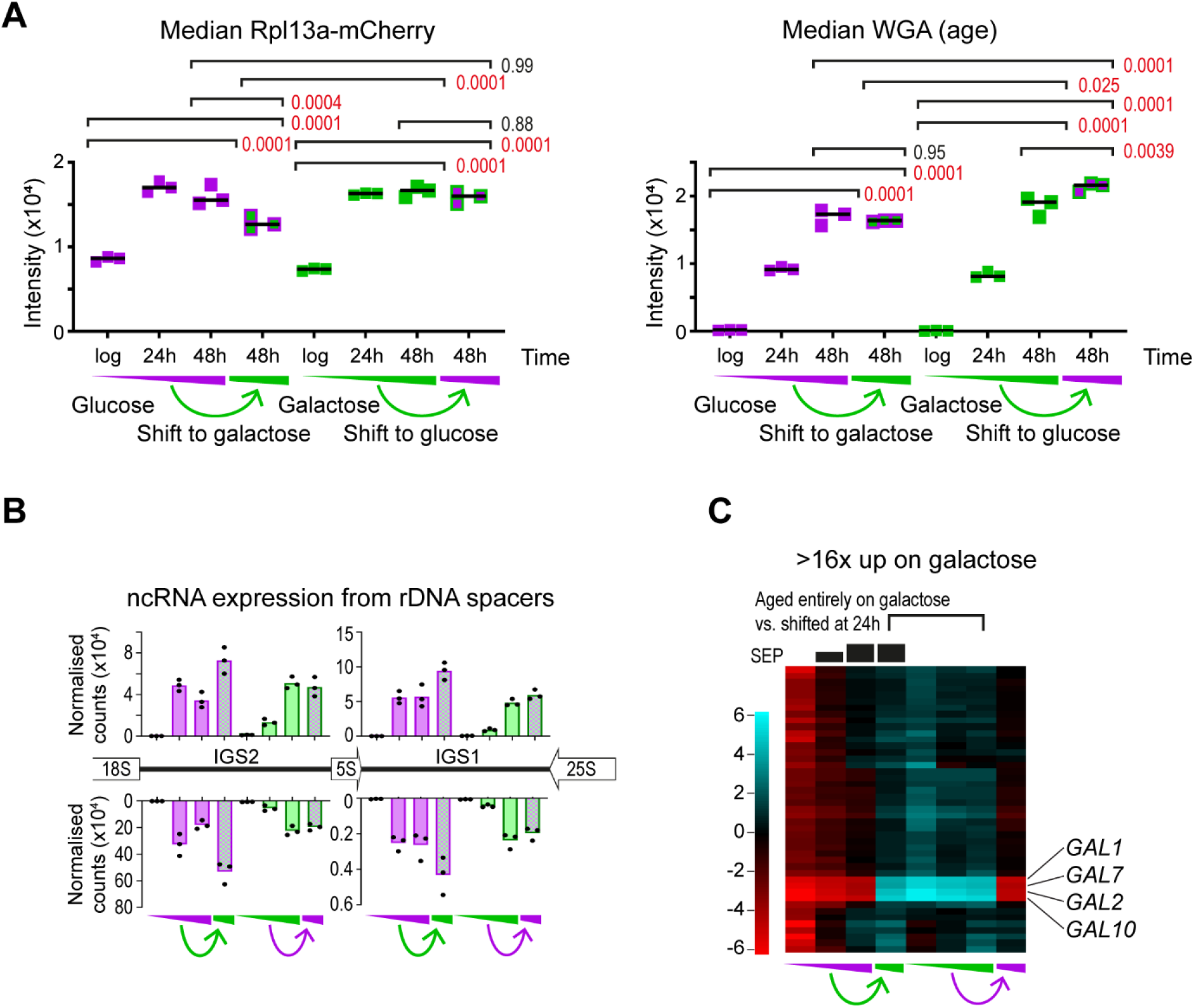
Supplement to media shifts determine the plasticity of ageing phenotypes. A: Rpl13a-mCherry and WGA intensity data for samples in Figure 6B. B: rDNA ncRNA abundances for media shift samples. C: Heatmap across all samples of genes either induced >16x on galactose relative to glucose at log phase, with primary galactose metabolic genes annotated. The extent of SEP entry is indicated diagrammatically above the plots based on Tom70-GFP intensities from B. Average of 3 biological replicates.

**Table S1:**
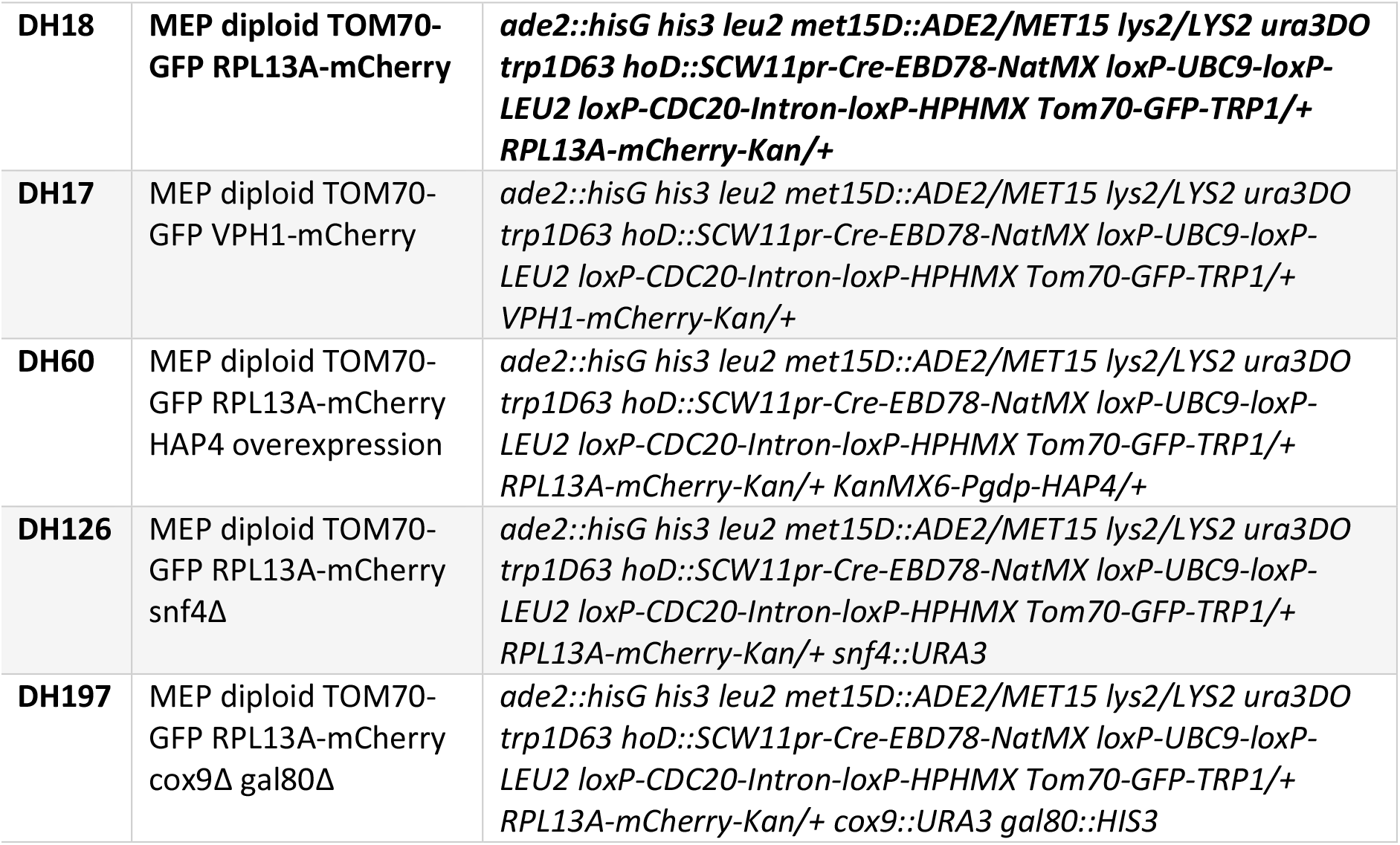
Strains used in this work. All strains are diploid derivatives of the MEP system (Lindstrom and Gottschling, 2009). TOM70-GFP, VPH1-mCherry and RPL13a-mCherry markers are heterozygous to avoid growth defect.

**Table S2:**
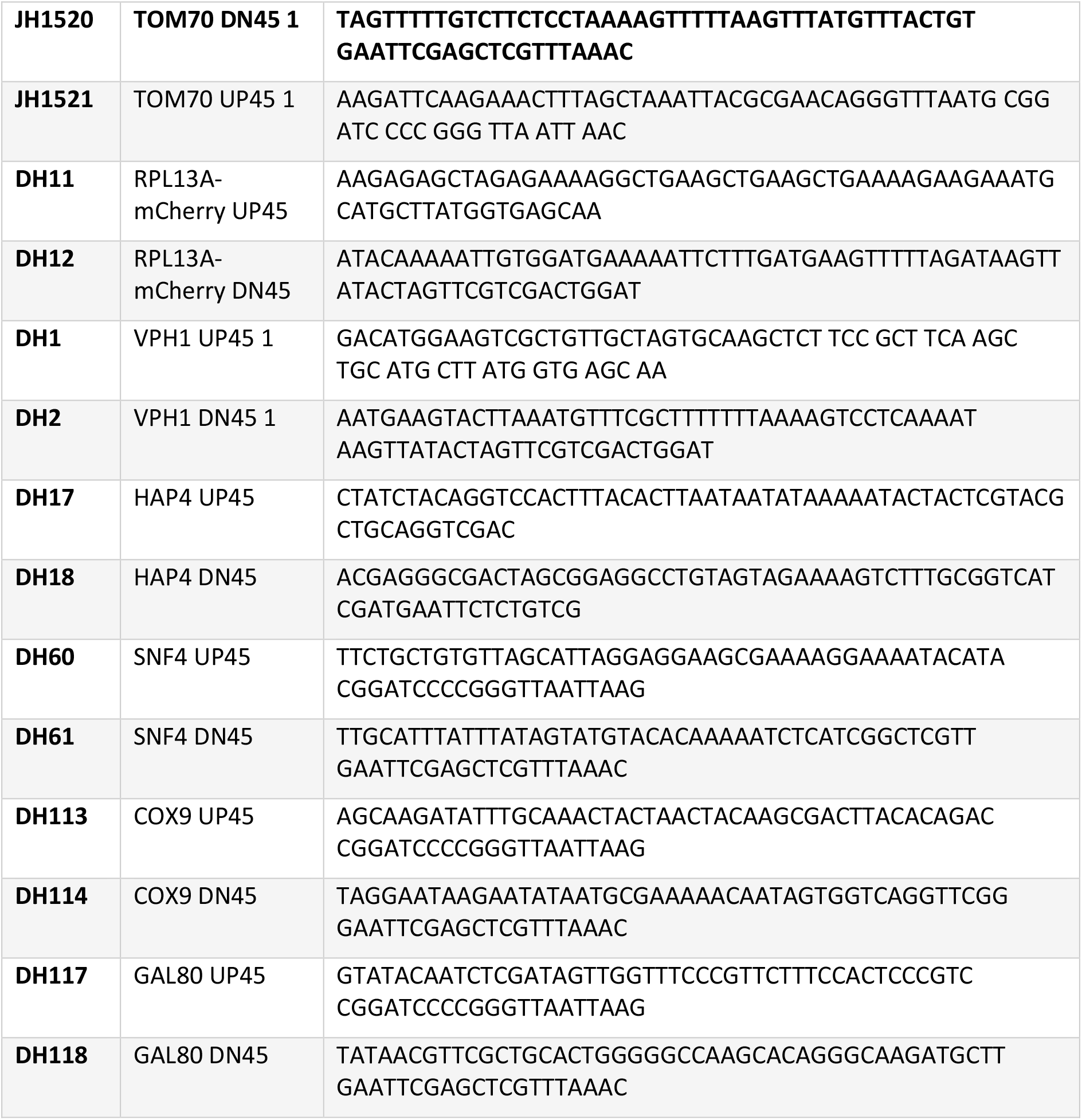
Oligonucleotides used for making strains.

